# Ocular delivery of different VCP inhibitory formulations prevents retinal degeneration in rhodopsin 255 isoleucine deletion mice

**DOI:** 10.1101/2025.06.25.661245

**Authors:** Bowen Cao, Regine Mühlfriedel, Merve Sen, Ana-Cristina Almansa-Garcia, Mathias W. Seeliger, Anne-Sophie Petremann-Dumé, Ellen Kilger, Anneli Vollert, Sylvia Bolz, Christine Henes, Paolo Caliceti, Stefano Salmaso, Marius Ueffing, Blanca Arango-Gonzalez

**Author notes:** Equal last authorship (Marius Ueffing, Blanca Arango-Gonzalez).

## Abstract

Rhodopsin-mediated autosomal dominant retinitis pigmentosa (*RHO*-adRP) is a progressive inherited retinal degenerative disorder currently lacking effective treatments. A recurrent 3-base pair deletion in the RHO gene, resulting in the loss of isoleucine at codon 255 or 256 (*RHO*^ΔI255^ or *RHO*^ΔI256^), has been identified in patients from the United Kingdom, Germany, Belgium, China, and Korea, suggesting a broad geographic distribution. This mutation leads to rhodopsin (RHO) misfolding, its retention in the endoplasmic reticulum (ER), and aggregation with wild-type (WT) RHO, ultimately triggering ER stress and photoreceptor degeneration. These aggregates are primarily cleared via the ER-associated degradation (ERAD) pathway, with valosin-containing protein (VCP) playing a key role in their retrotranslocation and proteasomal degradation. Pharmacological or genetic inhibition of VCP has shown neuroprotective effects in other models of adRP, but the poor aqueous solubility of VCP inhibitors and challenges in retinal drug delivery hinders clinical translation.

To overcome these limitations, we evaluated and compared three VCP-targeted therapeutic strategies in *Rho*^ΔI255^ knock-in mouse retinae: (1) small-molecule inhibitors (ML240, NMS-873) solubilized in DMSO, (2) ML240 encapsulated in monomethoxy-polyethylene glycol (mPEG)-cholane nanoparticles, and (3) small interfering RNA (siRNA) targeting VCP, delivered via magnetic nanoparticles. Neuroprotective effects were assessed in vitro in retinal explants and in vivo following intravitreal injection.

Our findings provide the first evidence that VCP inhibition restores RHO trafficking to the outer segments and prevents photoreceptor cell death in the *Rho*^ΔI255^ model. Among the three approaches, nanocarrier-encapsulated ML240 exhibited superior efficacy, enabling sustained drug delivery and enhanced photoreceptor protection. These results establish a preclinical proof-of-concept for nanocarrier-mediated VCP inhibition as a promising therapeutic strategy for *RHO*-adRP and potentially other ER-stress-related retinal degenerations.

**Graphical Abstract:** 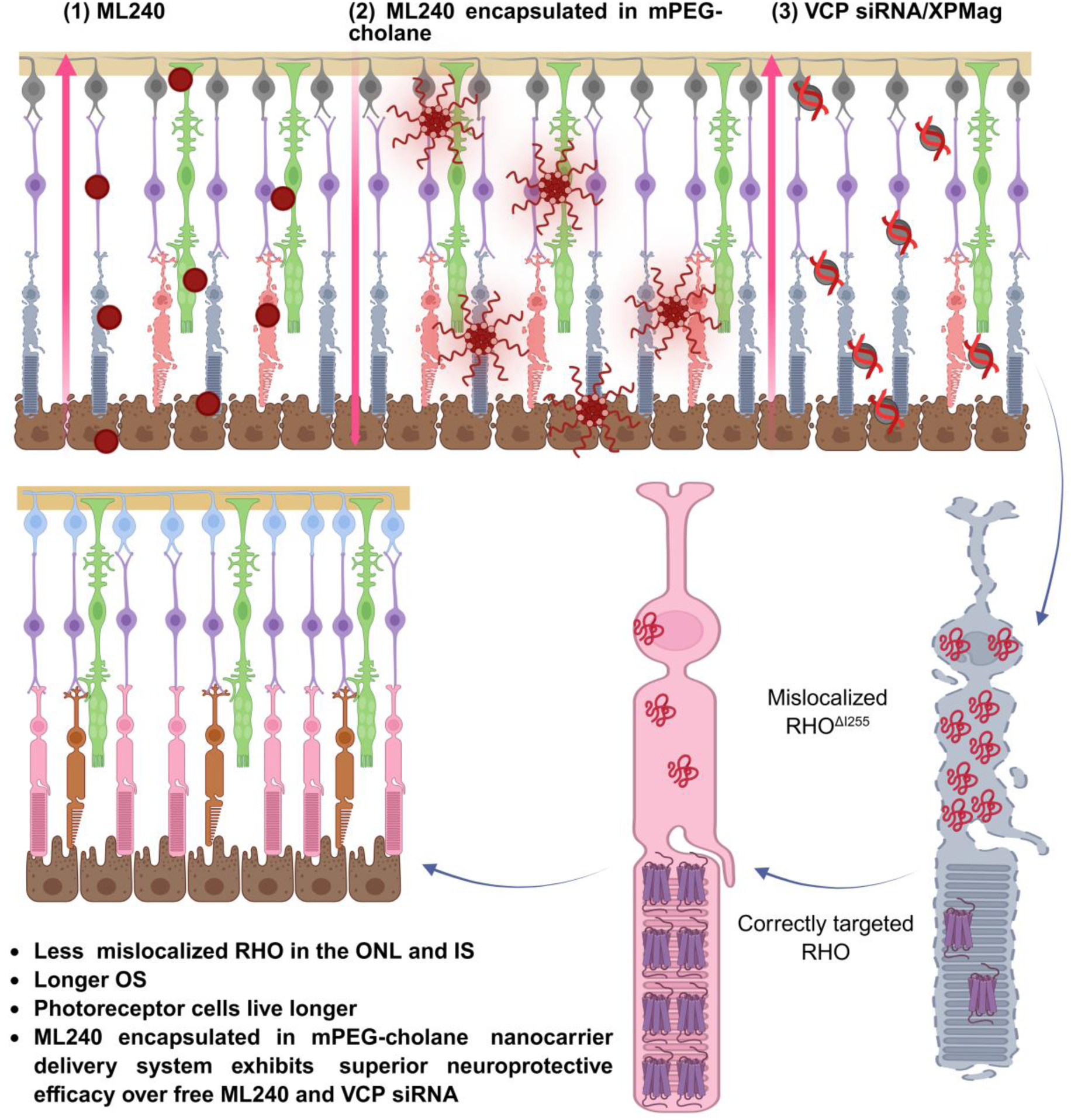

## Introduction

Retinitis pigmentosa (RP) is a heterogeneous group of inherited retinal degenerative disorders characterized by progressive degeneration of photoreceptor cells, ultimately leading to blindness. More than 200 mutations in the human rhodopsin gene (*RHO*), including point mutations, insertions, deletions, and complex rearrangements, have been associated with RP (Human Gene Mutation Database, HGMD). Among these, a 3-base pair (bp) deletion that removes one of the two isoleucines (I) at position 255 or 256 (*RHO*^ΔI255^/*RHO*^ΔI256^) was the first identified deletion mutation in *RHO*-associated autosomal dominant RP (*RHO*-adRP) [1]. This recurrent deletion mutation has been identified in multiple countries, including the United Kingdom, Germany, Belgium, China, and Korea, suggesting a broader geographic distribution beyond isolated populations [1–7]. Patients carrying this deletion mutation exhibit pronounced yet progressive retinal degeneration, including early-onset night blindness, gradual constriction of the visual field, and eventual irreversible vision loss [4].

Clinical observations have led to the classification of *RHO*-adRP patients into two major subtypes, Class A and Class B [8]. Patients with *RHO*^ΔI255^ exhibit characteristics of a Class B phenotype and are likely to fall into subclass B1. They present with milder photoreceptor degeneration and heterogeneous disease progression, which advances from the inferior to the superior retina. Notably, the P23H mutation (a proline-to-histidine substitution at codon 23), the most prevalent *RHO* mutation in North America [9], also falls within subclass B1.

Rhodopsin (RHO) is a G-protein-coupled receptor (GPCR) encoded by the *RHO* gene. It is synthesized in the rod inner segments (IS) and transported to the outer segments (OS), where it constitutes more than 90% of all proteins in the disk membrane [10]. Only correctly folded RHO can be transported to the rod OS and activated by photons to initiate the first step of phototransduction. Disruptions at any stage of this process, such as gene transcription, translation, protein folding, or trafficking to the designated location, can result in severe visual impairment [11].

The mutation on *RHO*^ΔI255^ has been reported to alter the protein folding process and is categorized as a class II adRP mutation, which is characterized by intracellular accumulation of misfolded proteins, endoplasmic reticulum (ER) retention, and defective RHO pigment formation [2, 12]. Our studies using cellular and retinal explant models have further revealed that RHO^ΔI255^ can sequester wild-type (WT) RHO at the ER, exerting a dominant-negative (DN) effect on the normal function of RHO^WT^, thereby subjecting both RHO variants to degradation via the valosin-containing protein (VCP)-dependent ER-associated degradation pathway (ERAD) [13]. Additionally, in the ΔI255 rhodopsin gene (*Rho*) knock-in mouse model, *Rho*^ΔI255^ triggers photoreceptor cell death by activating the apoptotic pathway dependent on caspase-3, as well as other cell death pathways involving calpain and poly (ADP-ribose) polymerase (PARP) [13]. Despite advances in understanding the pathogenesis and cell death mechanisms caused by *RHO*^ΔI255^, effective therapeutic strategies are still lacking.

Dysfunctional protein homeostasis is a hallmark of retinal degeneration. In our previous studies, we demonstrated that RHO^ΔI255^ aggregates are processed through the ERAD pathway, with VCP playing a key role in retrotranslocating the misfolded proteins from the ER to the cytoplasm for proteasomal degradation [13]. However, excessive accumulation of protein aggregates can overload ERAD capacity, disrupt ER homeostasis, and drive photoreceptor degeneration [14, 15]. Similar mechanisms have been reported for the misfolded RHO^P23H^ mutant, which induces excessive retrotranslocation activity, ultimately impairing ERAD function and triggering cell death [16, 17].

Pharmacological modulation of ERAD effectors has been explored as a potential treatment approach for retinal degeneration [16–21]. VCP is a known ERAD effector that facilitates the retrotranslocation of misfolded proteins from the ER to the proteasome for degradation [22]. Although counterintuitive, this protective effect may stem from reduced energy consumption associated with ERAD, redirection of misfolded protein aggregates towards autophagic degradation, and/or restoration of proteostasis through improved RHO trafficking [20].

Pharmacological inhibition of VCP by small molecule inhibitors or genetic VCP inactivation has been reported to significantly delay photoreceptor degeneration in *Rho*^P23H^ rodent models as well as in *Drosophila* expressing a misfolded RHO1^P37H^ protein [17, 20, 21, 23]. Pathological processes associated with *Rho*^P23H^ closely resemble those of *Rho*^ΔI255^, including the generation of ER-retained aggregates, impairment of the normal function of RHO^WT^, interaction with VCP, and degradation through the ERAD pathway [13, 16, 18]. Given the pathophysiological and clinical parallels between these two dominant *RHO* mutations, we hypothesized that treatment strategies effective against *Rho*^P23H^ might be beneficial for *Rho*^ΔI255^.

The primary goal of this study was to explore whether VCP inhibition using agents with distinct mechanisms of action can attenuate *Rho*^ΔI255^-associated retinal degeneration. Secondly, we aimed to identify an ocular drug delivery system (ODDS) that can be transferred to clinical practice. Effective drug administration to the posterior segment poses significant challenges owing to the unique anatomy and various barriers of the eye, including static (e.g., blood-aqueous and blood-retinal barriers) as well as dynamic barriers (e.g., conjunctival and choroidal blood flow, lymphatic clearance) [24]. In addition, the poor water solubility of VCP inhibitors further complicates drug delivery. Recently, various nanocarrier-based ODDS have been developed to improve bioavailability, stability, and sustained drug release [25]. Nano-based drug carriers, including nanocarriers based on amphiphilic polymers and nanoparticles, have shown promise in delivering therapeutic agents to both the anterior and posterior segments of the eye [26]. Among these, monomethoxy-polyethylene glycol (mPEG)-cholane-based nanocarriers have been successfully utilized to deliver biomacromolecules, including proteins [27], peptides [28], and small-molecule drugs [29, 30], yielding promising results in administering VCP inhibitors to the retina [31].

In addition to chemical drug applications, RNA-based therapeutics, particularly small interfering RNA (siRNA)-induced gene silencing, offer new possibilities for treating eye diseases, including RP [32–35]. However, efficient siRNA delivery to the retina remains challenging due to its susceptibility to enzymatic degradation, poor cellular uptake, and rapid clearance [36]. Magnetofection, an emerging technique for targeted gene delivery using magnetic nanoparticles (MNPs), has proven successful in overcoming these barriers [37–40]. A recent study in our lab provided evidence that VCP siRNA delivered via magnetofection can mitigate retinal degeneration in *Rho*^P23H^ rat models [41]. This study, therefore, assessed the feasibility of MNP-mediated siRNA delivery for VCP silencing in the *Rho*^ΔI255^ retina.

## Results

### 1. VCP inhibition protects photoreceptors from cell death in *Rho*^ΔI255/+^ mouse retinal explants

To investigate the neuroprotective potential of VCP inhibition in *Rho*^ΔI255/+^ retinal degeneration, we evaluated two small-molecule inhibitors, ML240 and NMS-873, which suppress VCP ATPase activity via distinct mechanisms. ML240 is a selective, reversible ATP-competitive inhibitor of VCP/p97 ATPase (IC50 = 110 nM), specifically targeting the D2-ATPase domain [42], whereas NMS-873 is a potent, selective, allosteric, non-ATP-competitive VCP/p97 ATPase inhibitor (IC50 = 30 nM), that binds to the D1 and D2 domains of adjacent subunits [43].

The effect of VCP inhibitors was tested using an organotypic retinal culture system comprising the neuroretina and retinal pigment epithelium (RPE), as previously described [44]. ML240 or NMS-873 were administered every 48 hours to the culture medium of *Rho*^ΔI255/+^ retinal explants. *Rho*^ΔI255/+^ retinae were isolated at PN14 and cultured for 6 days, as our previous study determined the peak of photoreceptor cell death in *Rho*^ΔI255/+^ mice to occur around postnatal day (PN) 20 [45]. ML240 and NMS-873 were initially dissolved in 0.5% DMSO and then applied to the culture medium to achieve concentrations of 20 μM for ML240 (which has previously demonstrated neuroprotection in *Rho*^P23H^ rat explants [20, 21]) and 0.5 μM NMS-873 (based on dose optimization results (**Fig. S1**)).

To quantitatively assess the potential neuroprotective effects of VCP inhibition, two parameters were measured: the percentage of TUNEL-positive cells (indicative of DNA fragmentation and thus of cell death) relative to the total number of cells in the outer nuclear layer (ONL) and the number of ONL cell rows.

ML240 treatment significantly reduced photoreceptor cell degeneration (**Fig. 1A**), as evidenced by a decrease in the percentage of TUNEL-positive dying cells (DMSO: 5.22 % ± 1.67, ML240: 2.28 % ± 0.89, *P* < 0.01. **Fig. 1B**), and corresponding preservation of the ONL structure, with a significant increase in photoreceptor cell rows (DMSO: 7.23 ± 0.52, ML240: 9.12 ± 0.84, *P* < 0.01. **Fig. 1C**).

**Figure 1.**
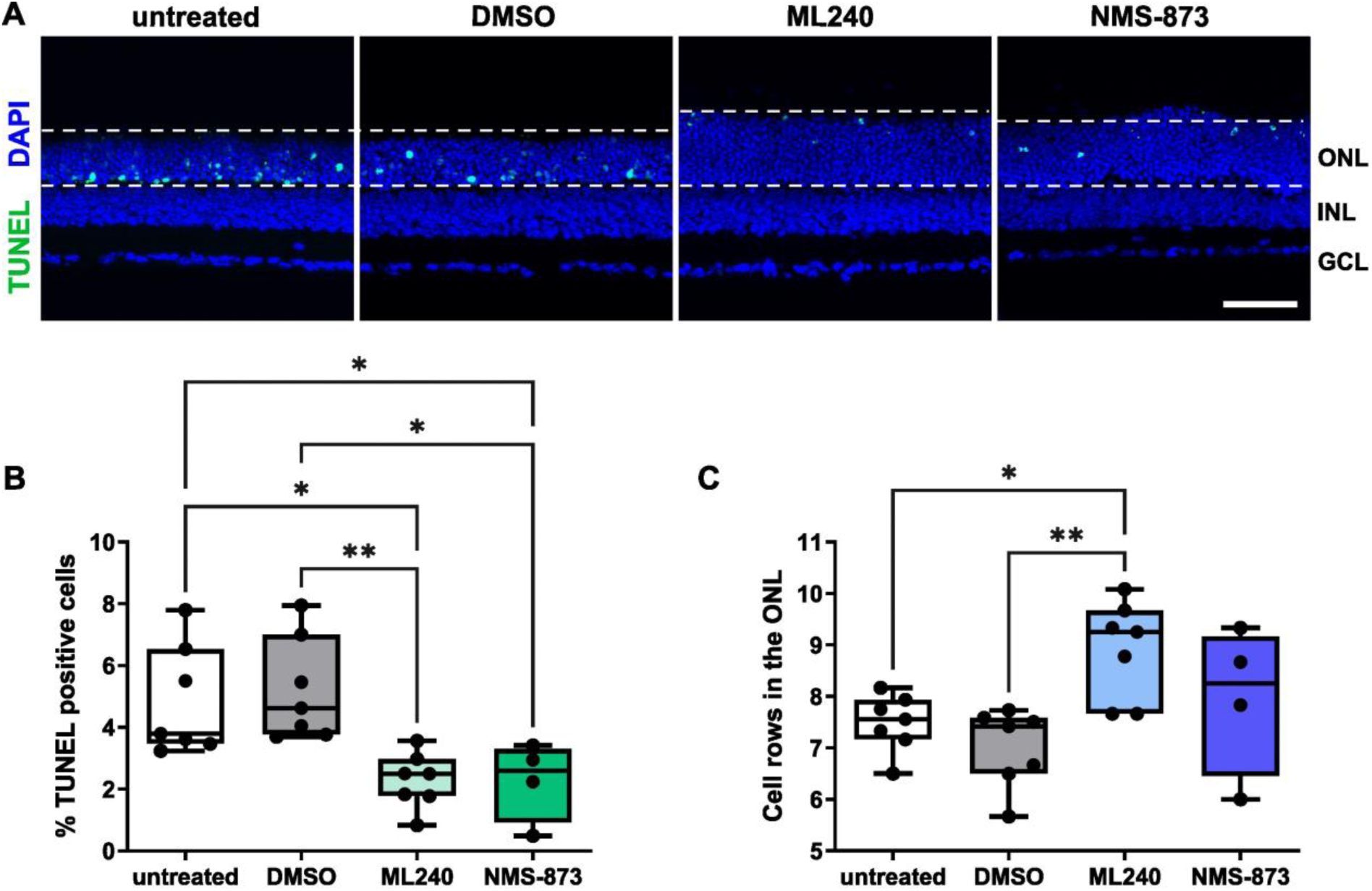
VCP inhibition significantly decreases photoreceptor cell death and promotes cell survival in *Rho*^ΔI255/+^ mouse retinae. RPE-attached retinae from *Rho*^ΔI255/+^ knock-in mice were isolated at postnatal day 14 (PN14), cultured for 6 days, and treated every second day with VCP inhibitors ML240 (20 μM) and NMS-873 (0.5 μM) dissolved in DMSO. Retinae without any treatment (untreated) or treated with 0.5% DMSO (DMSO) served as controls. **(A)** TUNEL assay reveals dying cells (green), while DAPI indicates all photoreceptor nuclei (blue). Scale bar: 50 μm. **(B)** Percentage of TUNEL-positive dying cells in the ONL was significantly reduced after treatment with either ML240 or NMS-873 compared to the DMSO control group. **(C)** Number of remaining cell rows in the ONL across groups. Retinae treated with ML240 showed significant preservation of the number of photoreceptor cell rows. Values were quantified from at least 4 retinae (black dots). Statistical analysis: box-and-whisker plots (median, quartiles, min-max); One-way ANOVA with Tukey’s multiple comparisons test; \**p* < 0.05, \*\**p* < 0.01. RPE: retinal pigment epithelium; OS: outer segment; IS: inner segment; ONL: outer nuclear layer; OPL: outer plexiform layer; INL: inner nuclear layer; IPL: inner plexiform layer; GCL: ganglion cell layer; NFL: retinal nerve fiber layer; HC: horizontal cell; BC: bipolar cell; AC: amacrine cell; GC: ganglion cell.

NMS-873 at the dosage of 0.5 μM significantly reduced photoreceptor apoptosis compared to vehicle control (DMSO: 5.22% ± 1.67, NMS-873: 2.27% ± 1.28, *P* < 0.05. **Fig. 1B**). Nevertheless, analysis of ONL architecture revealed that this dose did not significantly increase the number of photoreceptor nuclear rows (**Fig. 1C**). Higher concentrations of 1µM and 5 µM were associated with compromised retinal integrity, especially in the central retinal area (**Fig. S1D, G, J**).

### 2. VCP inhibitors restore RHO localization to the OS in *Rho*^ΔI255/+^ retinae

As a consequence of the *Rho*^ΔI255^ mutation, the RHO protein forms aggregates in the cytoplasm, which results in a marked reduction in OS length [2, 13, 45]. Impaired RHO trafficking and structural OS alterations precede photoreceptor cell death, suggesting that both protein aggregation and defective RHO transport play a pivotal role in disease progression [46]. Previous *in vitro* [18, 47] and *in vivo* [48] evidence has revealed that the aggregation of misfolded RHO is cytotoxic and contributes to cell death, suggesting that correcting RHO^ΔI255^ mislocalization may serve as a therapeutic strategy to delay retinal degeneration.

Given that VCP inhibition attenuates photoreceptor cell death in *Rho*^ΔI255/+^ retinae, we next sought to determine whether it also facilitates proper transportation of RHO to the OS.

In untreated and DMSO-treated control groups, overall RHO exhibited abnormal accumulation in the ONL and IS, with only a minor fraction properly trafficked to the OS (**Fig. 2A**), consistent with previous *in vivo* observations [45]. Following treatment with ML240 (20 μM) or NMS-873 (0.5 μΜ), a decrease in mislocalized RHO in the ONL and IS was observed, alongside restoration of RHO distribution within the OS, resembling a WT-like pattern (**Fig. 2A**).

**Figure 2.**
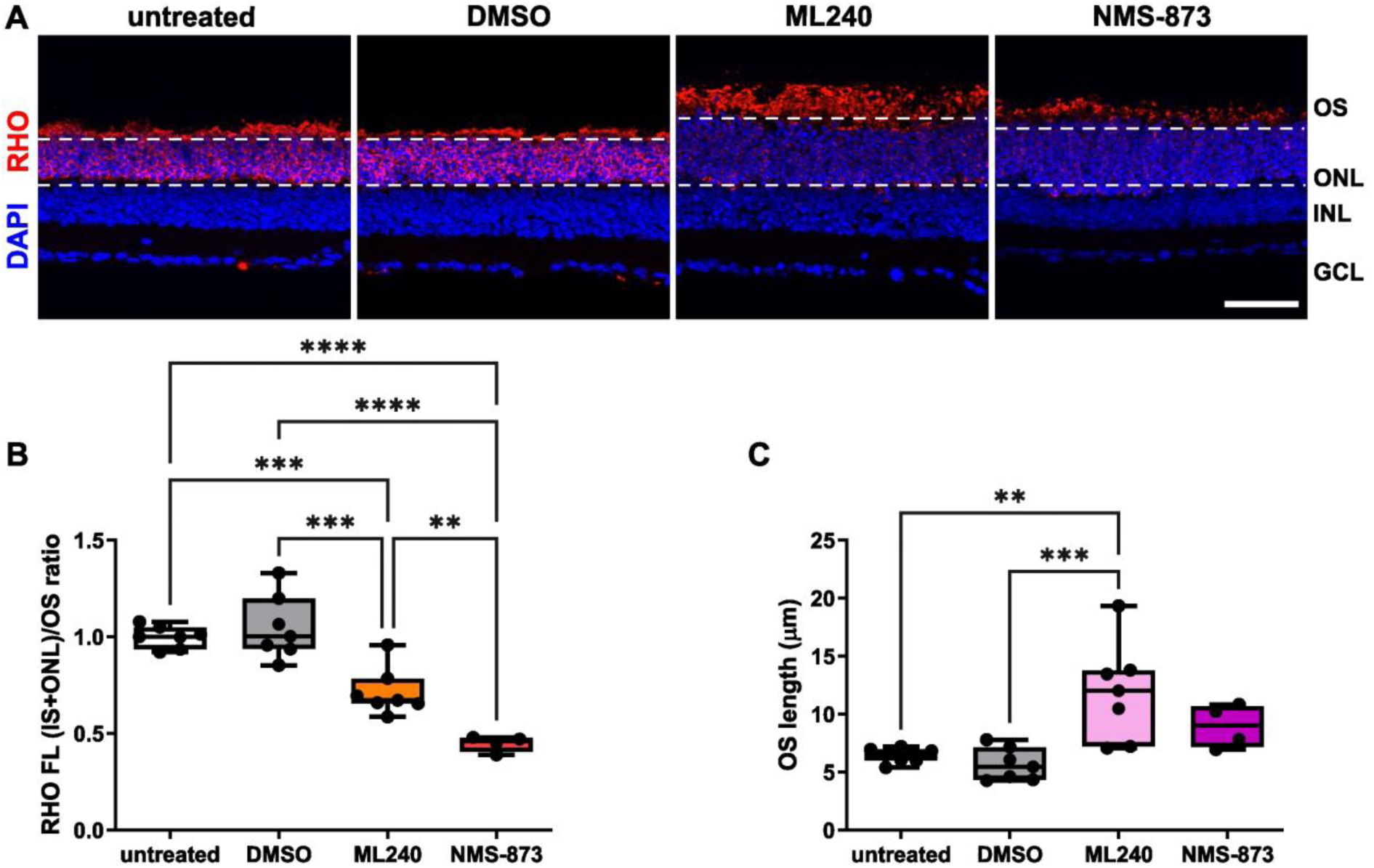
VCP inhibition reduces mislocalized RHO and restores the distribution of RHO to the OS. **(A)** RHO localization detected by immunofluorescent staining (red) in *Rho*^ΔI255/+^ retinal explants at PN20. Nuclei counterstained with DAPI (blue). RHO was aberrantly distributed in the ONL, and OS lengths were short in the untreated and DMSO-treated groups, while retinae treated with VCP inhibitors showed reduced RHO fluorescence in the ONL and better-preserved OS morphology. Scale bar: 50 μM. **(B)** The fluorescence intensity (FL) ratio between (IS+ONL) and OS was significantly decreased by VCP inhibition, suggesting a reduction of mislocalized RHO in ONL and IS. **(C)** Measurement of OS length in different groups. Retinae treated with ML240 significantly increased OS length compared to DMSO controls. Values were quantified from at least 4 retinae (black dots). Statistical analysis: box-and-whisker plots (median, quartiles, min-max); One-way ANOVA with Tukey’s multiple comparisons test; \*\**p* < 0.01, \*\*\**p* < 0.001, \*\*\*\**p* < 0.0001.

To further validate the beneficial effects of VCP inhibitors on RHO redistribution, the quantities of mislocalized and properly distributed RHO were quantified and compared. Mislocalized RHO is found in the IS and ONL and is represented by the ratio of the fluorescence intensity (FL) of IS + ONL to OS (IS+ONL)/OS, while the OS length is a measure of correctly distributed RHO. It should be noted that the IS in mutant retinae is challenging to identify. VCP inhibitors significantly decrease the (IS+ONL)/OS RHO FL ratio (DMSO: 1.05 ± 0.16, ML240: 0.72 ± 0.12, *P* < 0.001; NMS-873: 0.44 ± 0.04, *P* < 0.0001; **Fig. 2B**), indicating reduced retention of mutant RHO in ONL and IS compartments.

Additionally, VCP inhibition by ML240 significantly increased OS length, further supporting the role of VCP in promoting structural integrity and proper RHO trafficking (DMSO: 5.69 ± 1.38, ML240: 11.91 ± 4.24, *P* < 0.001. **Fig. 2C**). Additionally, OS length was marginally increased by NMS-873 at 0.5 μΜ, although this increase was not statistically significant (DMSO: 5.69 ± 1.38, NMS-873: 8.97 ± 1.86, *P* = 0.29. **Fig. 2C**).

These results demonstrate for the first time that VCP inhibition not only mitigates *Rho*^ΔI255^-associated photoreceptor degeneration but also restores physiological RHO distribution to the OS.

### 3. mPEG-cholane formulated ML240 efficiently promotes photoreceptor cell survival and restores proper RHO distribution at a lower dose in *Rho*^ΔI255/+^ retinal explants

Delivery and distribution of hydrophobic drugs like ML240 in the eye are hindered by poor aqueous solubility, limited tissue retention [20, 49], and potential cytotoxicity of solvents such as DMSO [50]. Therefore, to identify a VCP inhibitor formulation suitable for *in vivo* application in *Rho***^ΔI255^**^/+^ mice, a screening study was designed as a critical next step toward therapy development for human patients. Due to the narrow therapeutic window of NMS-873, where cytotoxicity was observed at doses only marginally higher than those with protective effects, the formulation was exclusively limited to ML240.

A nanocarrier system based on monomethoxy-polyethylene glycol (mPEG-cholane) was employed for this purpose. mPEG-cholane is an amphiphilic mPEG derivative that enhances drug solubility and provides sustained-release properties. The resulting formulation of mPEG-cholane nanoparticles encapsulating ML240 (mPEG-ML240) was exposed to *Rho*^ΔI255/+^ retinal explants at PN14. Explants were treated from the ganglion cell layer (GCL) side to mimic conditions close to *in vivo* intravitreal application (**Fig. 8F-b**) with either 2.5 μM mPEG-ML240 or the vehicle nanocarrier mPEG-cholane without ML240 (referred to as mPEG-vehicle) and incubated for 6 days. Media were replaced every 48 hours without additional drug supplementation. Untreated retinae were also included as a control. This experimental setup also enabled a direct comparison of the neuroprotective effect of mPEG-ML240 versus ML240 solubilized in DMSO. Again, four parameters were analyzed as indicators of neuroprotection for degenerative retinae: a. the percentage of TUNEL-positive cells, b. ONL cell rows, c. RHO fluorescence intensity ratio (IS+ONL)/OS, and d. OS length.

Although a 5 µM dose of mPEG-ML240 showed beneficial effects in the P23H model in previous work [31], a dose screening was conducted in the current mouse model. At this concentration, structural alterations were observed, including partial retinal swelling (data not shown). Administration of 2.5 µM mPEG-ML240 significantly reduced the percentage of TUNEL-positive cells compared to mPEG-vehicle alone (**Fig. 3A, B**) (mPEG-ML240: 1.86% ± 0.35 vs. mPEG-vehicle: 5.33% ± 1.16, *P* < 0.0001. A marked preservation of retinal structure was also detected, as indicated by a significant increase in the number of photoreceptor nuclear rows in the mPEG-ML240 group compared to mPEG-vehicle (mPEG-ML240: 9.07 ± 0.36 vs. mPEG-vehicle: 6.74 ± 0.39, *P* < 0.0001; **Fig. 3C**).

**Figure 3.**
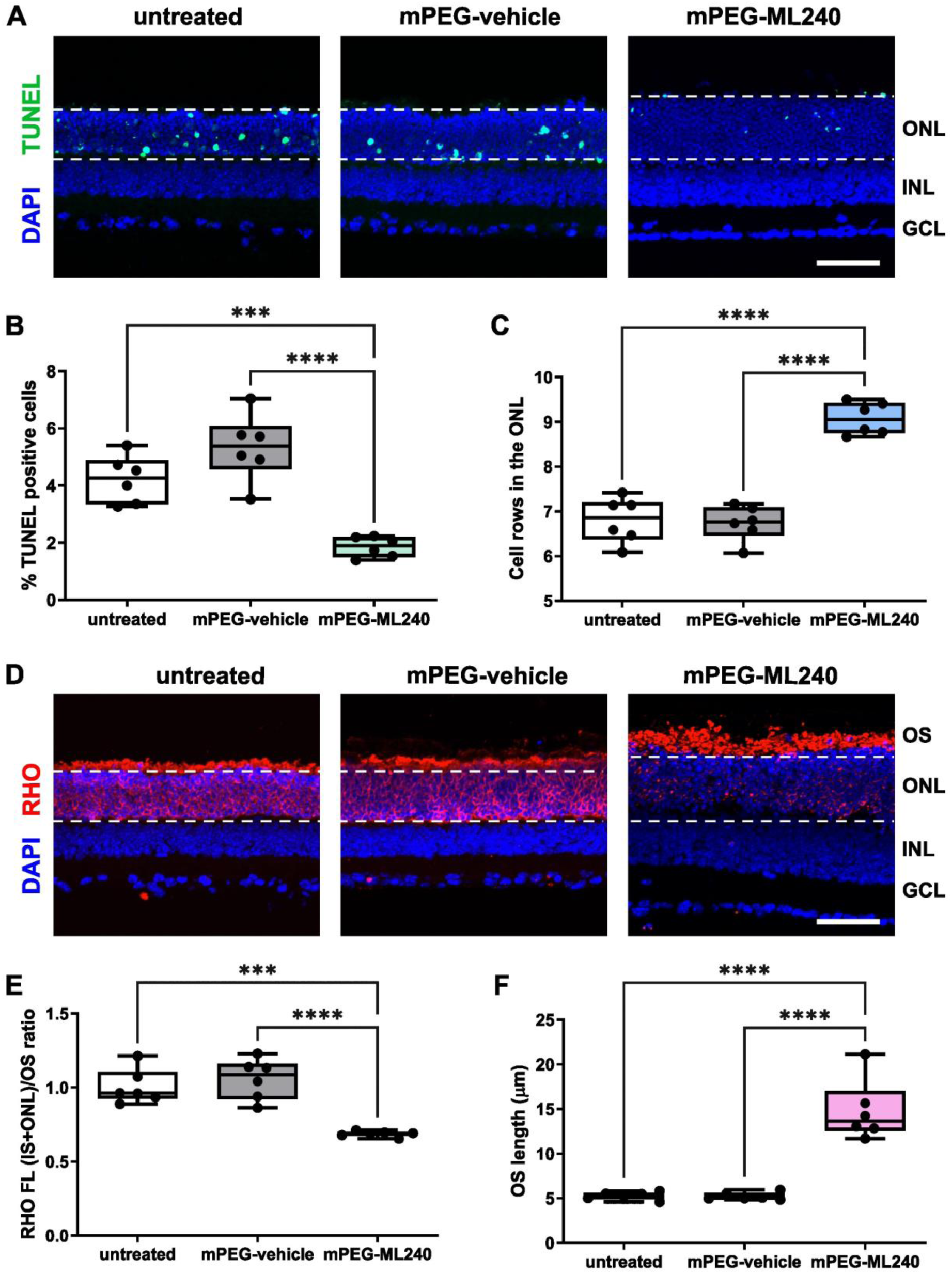
Delivery of ML240 by mPEG-cholane nanocarrier from the GCL side of the retina promotes photoreceptor cell survival and the correct trafficking of RHO in *Rho*^ΔI255/+^ retinae. (**A**) Retinae of *Rho*^ΔI255/+^ mice were explanted at PN14 and cultured for six days by administering the nanomicelles from the GCL side of the retina. Cryosections of retinae treated with mPEG-ML240 (ML240 concentration = 2.5 μΜ), mPEG-vehicle (vehicle concentration equivalent to that of mPEG-ML240 at 2.5 μΜ ML240), or untreated were stained with TUNEL assay to visualize dead cells (green), and DAPI nuclear counterstaining (blue). Scale bar: 50 μm. **(B)** Percentage of TUNEL-positive dying cells in the ONL. Compared to untreated and mPEG-vehicle groups, TUNEL-positive cells were significantly reduced after treatment with mPEG-ML240. **(C)** Quantification of the remaining ONL rows in different groups. Retinae treated with mPEG-ML240 showed a significant preservation of photoreceptor cell rows. **(D)** Fluorescent labeling (red) indicates RHO expression in *Rho*^ΔI255/+^ retinal explants. Nuclei were counterstained with DAPI (blue). RHO staining was mislocalized in the ONL in control groups, while retinae treated with mPEG-ML240 displayed a reduction of RHO in the ONL and better-preserved OS. Scale bar: 50μM. **(E)** Quantification of RHO FL ratio between (IS+ONL) and OS. mPEG-ML240 significantly decreased the FL ratio, suggesting a reduction of mislocalized RHO in the IS and ONL. **(F)** Measurement of OS length in different groups. Retinae treated with mPEG-ML240 significantly increased OS length compared to controls. Values were quantified from 6 retinae (black dots). Statistical analysis: box-and-whisker plots (median, quartiles, min-max); One-way ANOVA with Tukey’s multiple comparisons test; \*\*\**p* < 0.001, \*\*\*\**p* < 0.0001.

The mislocalization of RHO in the ONL and IS was also rescued by 2.5 μM mPEG-ML240 (**Fig. 3D**). Quantitative analysis of the FL ratio between (IS+ONL) and OS in the mPEG-ML240 group revealed a significant decrease in RHO mislocalization in the mPEG-ML240 group compared to both vehicle and untreated controls (mPEG-ML240: 0.69 ± 0.02, mPEG-vehicle: 1.06 ± 0.14, *P* < 0.0001; untreated: 1.01 ± 0.12, *P* < 0.001. **Fig. 3E**). Furthermore, mPEG-ML240 enhanced the correct targeting of RHO to the OS, as reflected by a significant increase in the OS length compared to the vehicle control group (mPEG-ML240: 14.78 ± 3.39, mPEG-vehicle: 5.23 ± 0.41, *P* < 0.0001; untreated: 5.27 ± 0.43, *P* < 0.0001. **Fig. 3F**).

Notably, the mPEG-vehicle exhibited no effect on either photoreceptor survival or cell death relative to untreated controls, supporting the biocompatibility and safety of the nanocarrier system. Additionally, unlike ML240 dissolved in DMSO, which required three administrations during the 6-day culture period, a single application of mPEG-ML240 produced a strong neuroprotective effect after only one application, achieving higher efficacy at a lower drug dose.

### 4. VCP silencing in *Rho*^ΔI255/+^ retinae by Reverse Magnetofection protects photoreceptors from degeneration and improves RHO trafficking

Pharmacological VCP inhibitors (ML240 or NMS-873) target only the ATPase activity of VCP but do not reduce its expression levels. To investigate whether reducing VCP protein expression could also be neuroprotective, small interfering RNA (siRNA) was employed to suppress VCP expression in *Rho*^ΔI255/+^ retinae.

VCP siRNA has been found to have potent gene-silencing effects in cell lines within 48-72 hours and in retinal explants for 72 hours [41]. Based on this timeline, we prepared *Rho*^ΔI255/+^ retinal explants at PN17 and conducted VCP silencing over three days until the peak of photoreceptor degeneration at PN20. Reverse Magnetofection was applied to retinal explants as described before [41], following the workflow illustrated in **Fig. 8E, F-c**. In brief, VCP siRNA (50 nM) was complexed with magnetic nanoparticles (XPMag) in serum-free medium (total volume of 100 μL) for 30 min, and the complexes were added to the culture medium. A magnetic plate was placed on top of the Transwell plate cover for 30 min to pull the complexes through all the retinal layers from the RPE to the GCL. The medium was replaced after 24 hours, and the explants were then incubated for an additional 2 days to evaluate the effect of VCP siRNA. It is worth mentioning that increasing the siRNA concentration to 100 nM did not improve the silencing efficiency or enhance photoreceptor protection (data not shown). As controls, retinal explants treated with XPMag only, XPMag/scrambled (Scr) siRNA, and untreated retinae were included.

VCP expression was first assessed by immunostaining (**Fig. 4A**). While VCP expression was observed across multiple retinal layers, we focused on the silencing efficiency within the ONL, where photoreceptors reside. Retinae treated with VCP siRNA showed a reduction in VCP immunofluorescent staining in the ONL (Fig. S2A). Quantification revealed a significant reduction of VCP expression in VCP siRNA-treated retinae compared to untreated and XPMag/scrambled control groups (VCP siRNA: 0.81 ± 0.19, untreated: 1.00 ± 0.09, *P* < 0.05; Scr siRNA: 1.04 ± 0.18, *P* < 0.05. **Fig. 4B**).

**Figure 4.**
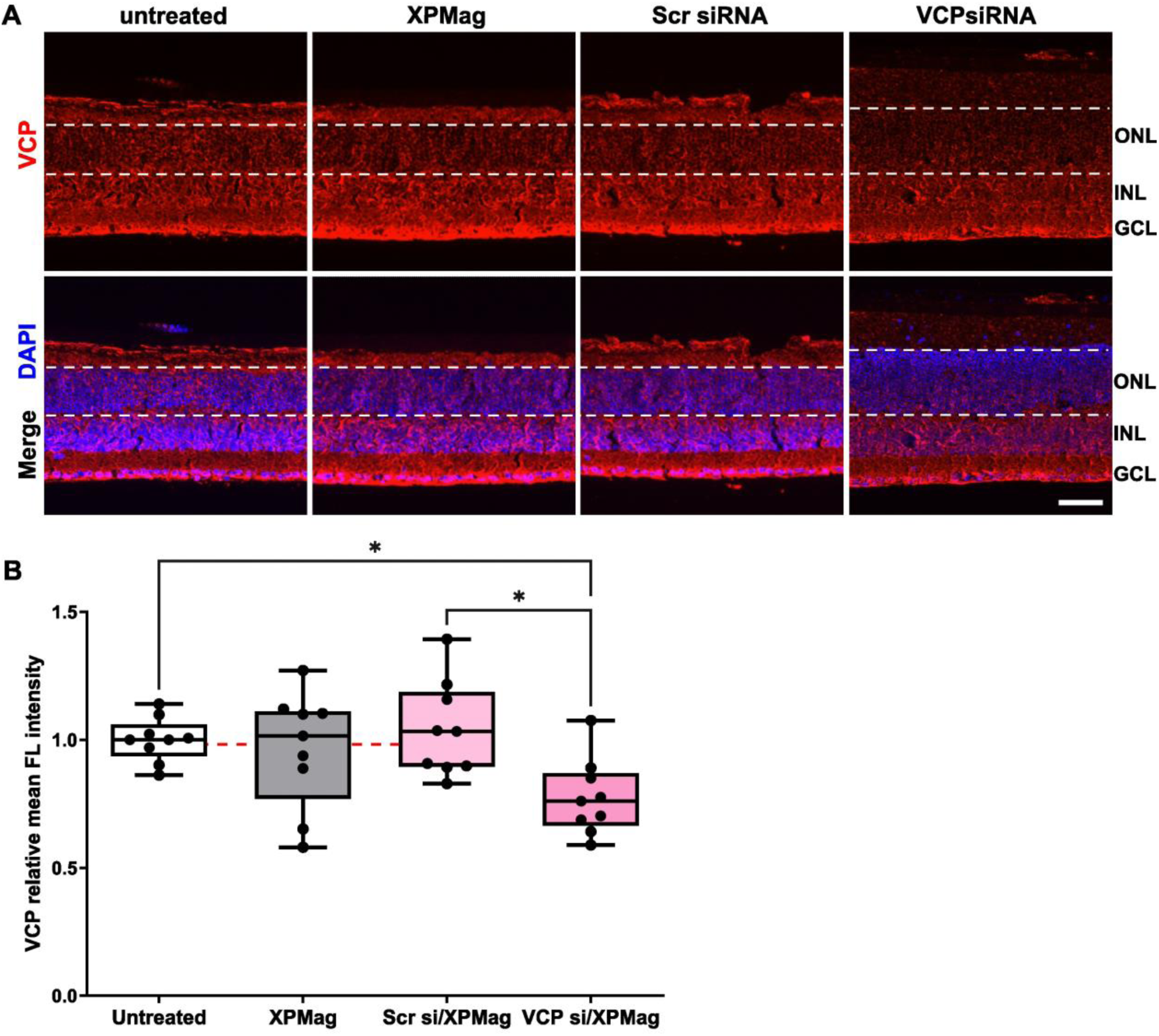
Gene silencing of VCP in *Rho*^ΔI255/+^ retinal explants by Reverse Magnetofection for 72. **h.** Retinae of *Rho*^ΔI255/+^ mice were explanted at PN17 and cultured for three days. Retinae were treated with 50 nM VCP siRNA/XPMag, 50 nM Scr siRNA/XPMag, XPMag alone, and medium only (untreated) by Reverse Magnetofection. (**A**) Explants were stained with specific VCP antibody (red) using nuclei counterstaining with DAPI (blue). Scale bar: 50 μm. (**B**) Quantification of VCP relative FL intensity in the ONL. The mean FL intensity was assessed by contouring the ONL regions. The relative value was calculated by comparing the values of other three groups with those of the untreated group. VCP siRNA significantly reduced VCP expression in the ONL compared with untreated and XPMag/scrambled controls. The mean VCP FL intensity in the VCP siRNA group was lower than that in the XPMag-treated retinae, but this was not significant. The red dashed line indicates the mean value of VCP FL intensity in the XPMag group. Values were quantified from 9 retinae (black dots). Statistical analysis: box-and-whisker plots (median, quartiles, min-max); One-way ANOVA with Tukey’s multiple comparisons test; \**p* < 0.05.

To evaluate the safety and efficacy of Reverse Magnetofection and the neuroprotective effect of VCP suppression, photoreceptor cell death (via TUNEL staining) and ONL integrity (via DAPI staining) were analyzed (**Fig. 5A**). The percentage of TUNEL-positive cells in untreated retinae was 2.01 % ± 0.59. Applying XPMag alone or XPMag/Scr siRNA did not cause a significant increase in cell death, suggesting that Reverse Magnetofection was safe to apply.

**Figure 5.**
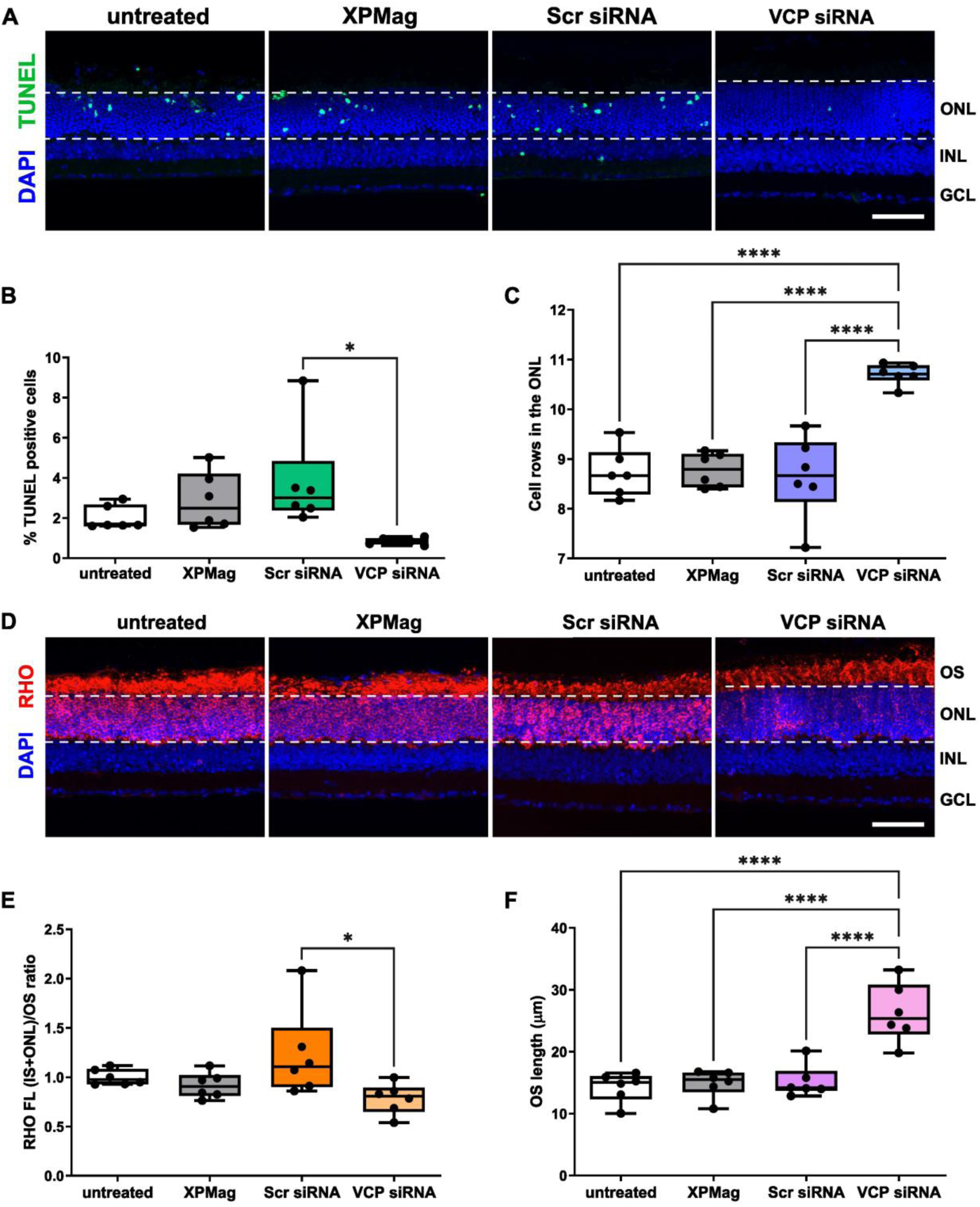
VCP silencing via Reverse Magnetofection is neuroprotective in *Rho*^ΔI255/+^ retinae. **(A)** TUNEL assay in *Rho*^ΔI255/+^ **r**etinal cryosections (green), with nuclear counterstaining using DAPI (blue). Scale bar: 50 μm. **(B)** Percentage of TUNEL-positive dying cells in the ONL. The percentage of TUNEL-positive cells was significantly reduced in the VCP siRNA group compared to the scrambled siRNA (Scr siRNA) control group. **(C)** Quantification of the number of remaining ONL cell rows. VCP silencing significantly preserved photoreceptor cell structure. **(D)** RHO expression (red) in *Rho*^ΔI255/+^ retinal explants. Nuclei were counterstained with DAPI (blue). RHO is predominantly mislocalized in the ONL in control groups, while it was mainly localized in the OS in VCP siRNA-treated retinae. Scale bar: 50μM. **(E)** Quantification of RHO FL ratio between (IS+ONL) and OS. VCP siRNA significantly decreased the FL ratio, suggesting a reduction of mislocalized RHO in the IS and ONL. **(F)** Measurement of OS length in different groups. VCP-silenced retinae exhibited significantly increased OS length compared to controls. Values were quantified from at least 6 retinae (black dots). Statistical analysis: box-and-whisker plots (median, quartiles, min-max); One-way ANOVA with Tukey’s multiple comparisons test; \**p* < 0.05, \*\*\*\**p* < 0.0001.

Retinae treated with VCP siRNA showed a significant reduction in TUNEL-positive cells (VCP siRNA: 0.84 % ± 0.16, Scr siRNA: 3.82 ± 2.52, *P* < 0.01; **Fig. 5B**), indicating a protective effect on photoreceptors. Furthermore, VCP silencing significantly increased the number of ONL nuclear rows compared to scrambled controls (VCP siRNA: 10.7 ± 0.21, Scr siRNA: 8.65 ± 0.84, *P* < 0.0001; **Fig. 5C**), reflecting improved photoreceptor survival.

To determine the effect of VCP silencing on RHO localization, RHO mislocalization and trafficking to the OS was evaluated (**Fig. 5D**). We saw a significant reduction in RHO mislocalization, reflected by a lower FL ratio compared to scrambled siRNA controls (VCP siRNA: 1.23 ± 0.48, Scr siRNA: 0.78 ± 0.16, *P* < 0.05. **Fig. 5E**). In addition, reduced VCP expression led to enhanced delivery of RHO to the OS, as indicated by a significant increase in OS length in VCP-silenced retinae compared to all control group (VCP siRNA: 26.27 ± 4.77, XPMag: 15.20 ± 2.59, *P* < 0.0001; Scr siRNA: 14.92 ± 2.20, *P* < 0.0001; untreated: 14.28 ± 2.38, *P* < 0.0001. **Fig. 5F**).

These results demonstrate that VCP suppression by siRNA protects photoreceptors from degeneration and restores RHO trafficking to the OS in *Rho*^ΔI255/+^ retinae without inducing toxicity.

### 5. Comparative analysis of the neuroprotective effect of three VCP inhibition strategies

In this study, three different VCP inhibition strategies were employed, including free small molecule drugs (ML240 or NMS-873 dissolved in DMSO), nanoparticles (mPEG-cholane-encapsulated ML240), and gene silencing via magnetic nanoparticles (XPMag/VCP siRNA complexes). Although all three formulations conferred measurable neuroprotective effects on *Rho*^ΔI255/+^ retinae, their efficacy varied across key outcome metrics, including photoreceptor cell survival, RHO trafficking, and OS structural preservation.

To enable direct comparison, data were normalized by calculating the relative values of treated explants to their respective untreated controls, and these ratios were compared across the three VCP inhibition groups.

Among the formulations, mPEG-ML240 consistently outperformed the other treatments in increasing the number of photoreceptor cell rows (**Fig. 6B**) and restoring proper RHO distribution to the OS (**Fig. 6D**), compared to free ML240 and VCP siRNA treatments. Notably, mPEG-ML240 showed only a slight advantage over the other two treatments in reducing the percentage of TUNEL-positive cells (mPEG-ML240: 0.44 ± 0.09, ML240: 0.49 ± 0.21, VCP siRNA: 0.46 ± 0.21. **Fig. 6A**). Reconstituting proper RHO trafficking, mPEG-ML240 performed far better than VCP siRNA, but was not significantly superior to ML240 alone (mPEG-ML240: 0.68 ± 0.02, VCP siRNA: 0.95 ± 0.19, *P* < 0.05. **Fig. 6C**).

**Figure 6.**
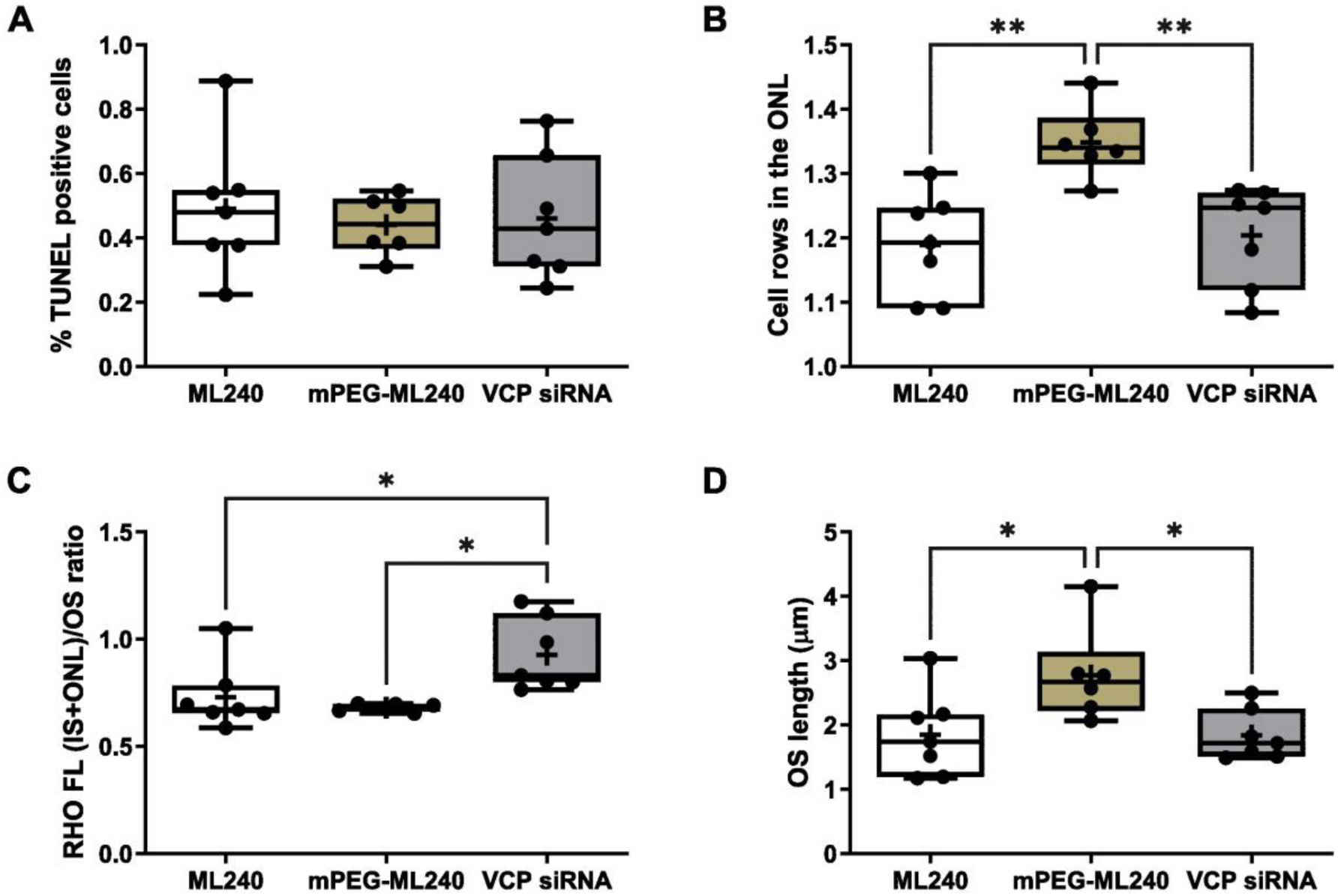
Comparison of neuroprotective parameters in *Rho*^ΔI255/+^ retinae treated with different VCP inhibition strategies. Analysis of percentage of TUNEL-positive cells **(A)**, ONL cell rows **(B)**, ratio of RHO FL between (IS+ONL) and OS **(C)**, and OS length **(D)** in treated retinae relative to untreated retinae. Values were quantified from at least 6 retinae (black dots). Data presented as mean ± SD; Statistical analysis: box-and-whisker plots (median, quartiles, min-max); One-way ANOVA with Tukey’s multiple comparisons test; \**p* < 0.05, \*\**p* < 0.01, + indicates the mean value.

Overall, mPEG-ML240 demonstrated the most favorable profile, combining enhanced efficacy with a lower required dose and prolonged bioavailability.

### 6. Intravitreal injection of mPEG-ML240 provides a long-term neuroprotective effect in Rho^ΔI255/+^ mice

As shown above, mPEG-ML240 can achieve a neuroprotective effect for 6 days in retinal explants of *Rho*^ΔI255/+^ mice. To determine whether mPEG-ML240 also provides neuroprotection *in vivo* over a longer period, intravitreal injections were performed in *Rho*^ΔI255/+^ mice, and retinal tissue was analyzed 16 days later.

Higher doses of mPEG-ML240 were tested *in vivo,* as intravitreally applied drugs can be cleared from the eye via the flow of aqueous humor and the ocular vasculature [49]. Accordingly, *Rho*^ΔI255/+^ mice received a single treatment of 5 µM mPEG-ML240. As a control, the other eye of the same mouse was treated with the same concentration of mPEG-cholane alone (vehicle group). All mice were injected at PN14 and kept for 16 days until PN30. Considering the superior-inferior hemispheric asymmetry of photoreceptor degeneration in *Rho*^ΔI255/+^ retinae [45], we analyzed and compared four retinal sections: inferior peripheral, inferior middle-peripheral, superior middle-peripheral, and superior peripheral.

*Rho*^ΔI255/+^ retinae treated with 5 µM mPEG-ML240 demonstrated decreased cell death in the whole retinal section (**Fig. 7A**), with notable reductions in inferior middle-peripheral and superior middle-peripheral areas as compared to vehicle-treated control (% TUNEL positive cells in inferior middle-peripheral region treated with mPEG-ML240: 0.71 ± 0.49, vehicle: 1.50 ± 0.50, *P* < 0.05; superior middle-peripheral treated with mPEG-ML240: 0.58 ± 0.29, vehicle: 0.97 ± 0.29, *P* < 0.05. **Fig. 7B**). In addition, retinae treated with mPEG-ML240 displayed an increase of ONL cell rows in the superior peripheral region (cell rows in superior peripheral area treated with mPEG-ML240: 8.38 ± 0.59, vehicle: 7.32 ± 0.93, *P* < 0.05. **Fig. 7C**. In other retinal regions, the number of ONL rows was slightly increased compared to control eyes, but this difference was not statistically significant.

**Figure 7.**
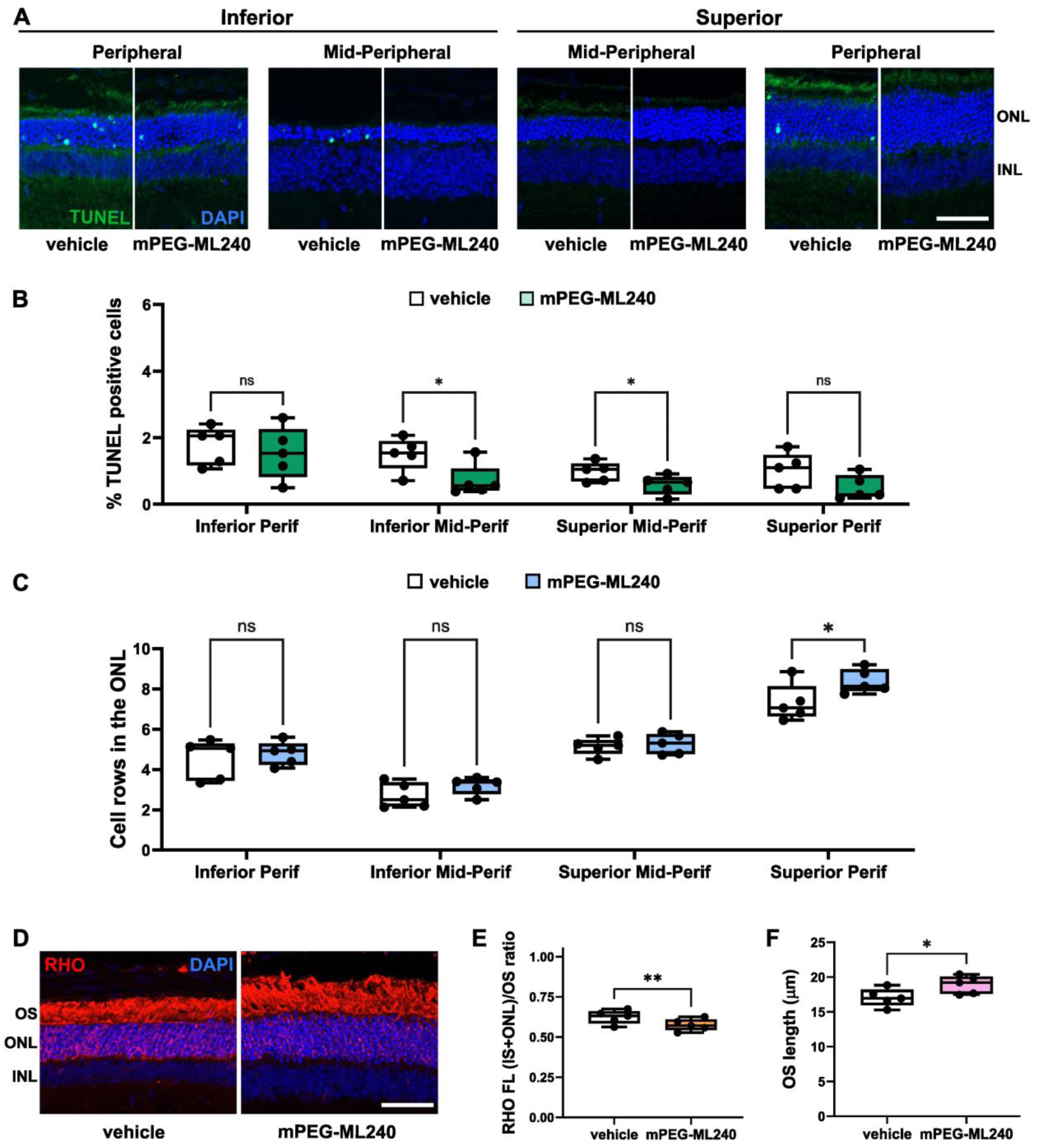
A single intravitreal injection of mPEG-ML240 exerts long-term neuroprotective effects on *Rho*^ΔI255/+^ retinae. **(A)** Retinal cross-sections from PN30 were stained using TUNEL assay (green) and DAPI nuclear counterstaining (blue). Four different regions from the superior-inferior hemispheres were examined, including inferior peripheral (Inferior Perif), inferior middle-peripheral (Inferior Mid-Perif), superior middle-peripheral (Superior Mid-Perif), and superior peripheral (Superior Perif). Scale bar: 50µm. **(B)** Percentage of TUNEL-positive cells in the ONL. Treatment of *Rho*^ΔI255/+^ retinae with 5 µM mPEG-ML240 significantly reduced cell death in the Mid-Perif regions of both superior and inferior retinae compared with vehicle controls. **(C)** Analysis of cell rows in the ONL of vehicle- and mPEG- ML240-treated retinae. The ONL cell rows increased in the retinae treated with 5 µM mPEG-ML240, with a significant increase in the Superior Perif area. **(D)** Fluorescent labeling of RHO (red) in vehicle- and 5 µM mPEG-ML240-treated *Rho*^ΔI255/+^ retinae. Nuclei were counterstained with DAPI (blue). In *Rho*^ΔI255/+^ retinae treated with mPEG-ML240, RHO was partially restored to the OS. Scale bar: 50μM. Quantification of RHO FL ratio between (IS+ONL) and OS shown in **E,** and measurement of OS length shown in **F**. Values were quantified from 5 retinae (black dots). Statistical analysis: box-and-whisker plots (median, quartiles, min-max); paired two-tailed *t*-test; \**p* < 0.05, \*\**p* < 0.01, ns: no significance.

**Figure 8.**
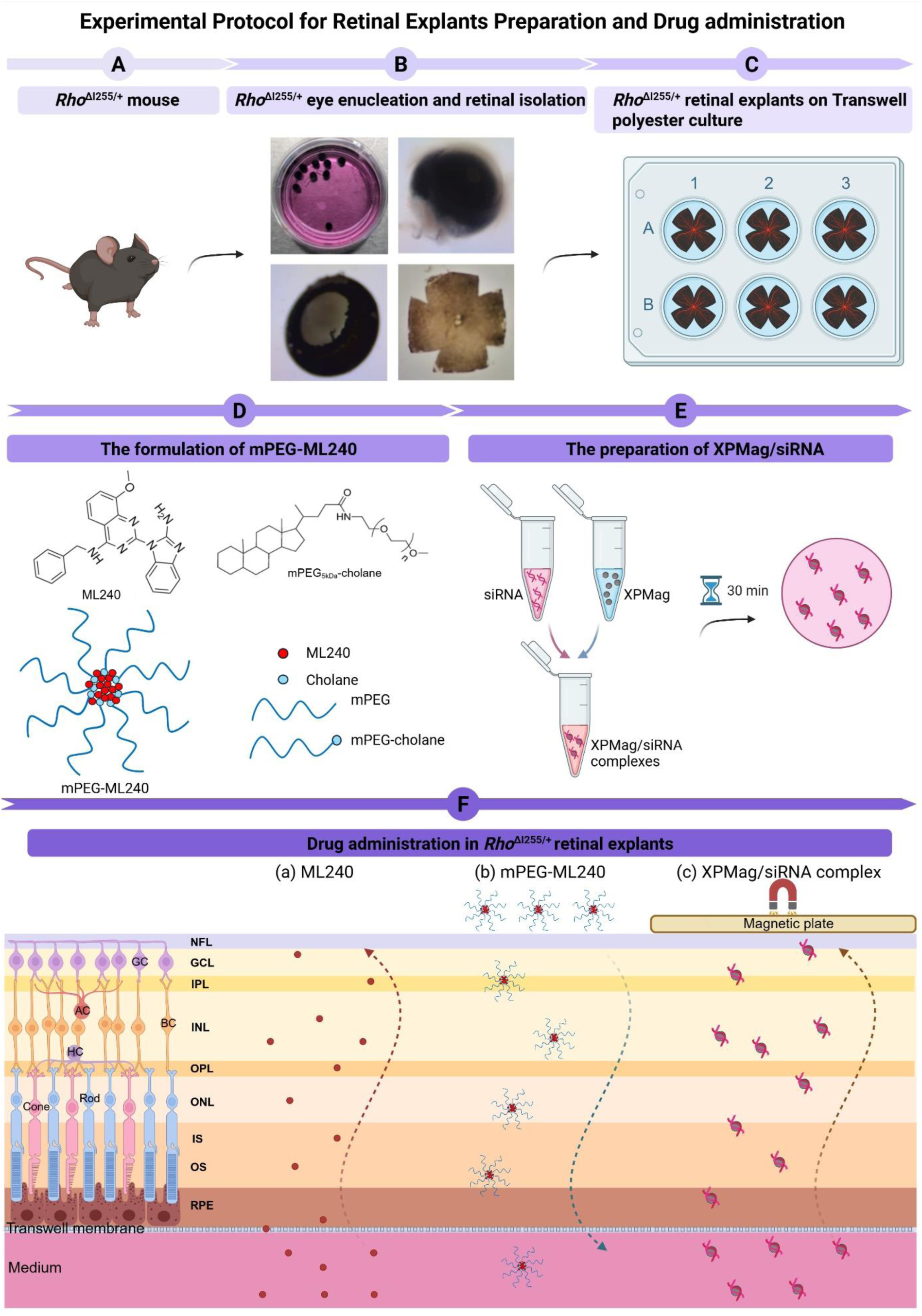
Schematic illustration of retinal explant preparation and drug administration. (**A**) *Rho*^ΔI255/+^ mice. (**B**) Preparation of *Rho*^ΔI255/+^ retinal explants. The eyes were enucleated in an aseptic environment. Sclera, choroid, and lens were carefully removed, leaving only neural retina and RPE. Four radical incisions were cut to flatten the retina. (**C**) The entire retina with its adherent RPE was mounted flat polycarbonate Transwell® membranes, with the GCL facing up. (**D**) Chemical structures of ML240, mPEG_5kDa_-cholane, and the self-assembled ML240 encapsulated nanoparticle system. (**E**) Preparation of Reverse Magnetofection for transfection of siRNA. Briefly, siRNA was added into magnetofection nanoparticle (XPMag), incubated for 30 min to obtain XPMag/siRNA complexes, and then the complexes were added to the retinal culture medium. (F) Administration of three VCP inhibitory formulations into retinal explants. ML240 dissolved into DMSO was administrated into the culture medium and retinae were treated for 6 days (**a**), mPEG-ML240 was applied on the GCL side of the retina for 6 days (**b**), and XPMag/siRNA complexes were applied into the medium, attracted by a magnetic plate for 1 h, and then incubated the transfected retinae for 72 h (**c**). Created with BioRender.com.

5 µM of mPEG-ML240 also significantly reduced the mislocalization of RHO in both ONL and IS (**Fig. 7D)**, as indicated by the decrease in the ratio of RHO fluorescence intensity between (ONL+IS) and OS (mPEG-ML240: 0.58 ± 0.04, vehicle: 0.62 ± 0.04, *P* < 0.01; **Fig. 7E**). Immunofluorescence staining also pointed to increased OS lengths following mPEG-ML240 treatment; however, quantification of differences did not reach statistical significance (mPEG-ML240: 18.92 ± 1.29, vehicle: 17.06 ± 1.30, *P* < 0.01; **Fig. 7F**).

These results indicate a long-term neuroprotective effect following a single intravitreal injection of mPEG-cholane-encapsulated ML240 in *Rho*^ΔI255/+^ mice, supporting sustained drug activity potentially due to prolonged release from the nanocarrier system.

## Discussion

The *RHO*^ΔI255^ mutation has been the first identified deletion mutation in the *RHO* gene that causes *RHO*-associated autosomal dominant retinitis pigmentosa (*RHO*-adRP) in Europe [1, 4, 51]. It has also been reported in Asia, including China [1, 4, 6, 7, 51, 52]. Previous studies [2, 12], as well as our recent data [13], indicate that this class II adRP mutation produces ER-associated protein aggregates targeted for degradation via the ER-associated degradation (ERAD) pathway. VCP, a key ERAD effector, mediates the retrotranslocation of these aggregates for proteasomal clearance. Overactivation of this pathway has been linked to ER stress and photoreceptor degeneration [17]. Although the pathogenicity of this mutation is well-established, its therapeutic implications remain largely unexplored.

This study, therefore, analyzed whether inhibiting VCP could attenuate retinal degeneration in *Rho*^ΔI255/+^ mice by limiting overactive ERAD. The *Rho*^ΔI255/+^ mouse model replicates the human disease phenotype [45], exhibiting progressive photoreceptor loss and diminished production of functional RHO in the OS. Treatment with ML240 or NMS-873 decreased photoreceptor cell death and improved RHO localization. These effects parallel findings in *Rho*^P23H^ models [17, 20, 21], another class II adRP mutation. Mutant RHO^P23H^ exhibits similar pathological mechanisms to RHO^ΔI255^, exerting a toxic dominant misfunction, also known as gain-of-function (GOF) in other publications, and dominant-negative effects [13, 53]. In cultured cells, RHO^ΔI255^, as well as RHO^P23H^, are retained in the ER, inducing the unfolded protein response (UPR) and ultimately leading to apoptosis [19, 46, 54]. Furthermore, both mutant aggregates are polyubiquitinated, retrotranslocated from the ER to the cytosol by cellular chaperone VCP, and degraded by the VCP-dependent ERAD machinery [13, 16]. Whether these parallels translate into similar therapeutic windows remains to be determined.

The role of VCP in retinal degeneration appears counterintuitive: one would expect that VCP activity facilitates the clearance of protein aggregates from the cell, thereby preventing their pathological accumulation. Consequently, inhibition of VCP would be expected to promote the accumulation of misfolded proteins in the ER and within the cytosol. In contrast, VCP inhibition reduced RHO accumulation in the ONL and IS, prevented aggregate formation, and enhanced trafficking to the OS. Efficient RHO transfer from the ONL and IS to the OS following VCP inhibition correlated with reduced photoreceptor cell death and structural degeneration. However, it remains unclear when and how VCP inhibition restores RHO localization to the OS and confers photoreceptor protection, underscoring the need for further investigation.

Excessive accumulation and aggregation of poly-ubiquitinated proteins have been reported to interfere with their clearance via the ubiquitin-proteasome system (UPS) or via autophagy [18, 47]. This interference can lead to a buildup of unprocessed proteins, which further interfere with the proper processing of WT or other proteins. As RHO proteins form dimers [13, 55], misfolded RHO^ΔI255^ likely binds WT RHO, thereby preventing the trafficking of RHO^WT^ from reaching the plasma membrane as well as the outer segment. This likely causes its dominant-negative effect, resulting in ER stress, induction of UPR signaling, and activation of pro-apoptotic signaling [19, 56]. Under normal physiological conditions, activation of PERK, IRE1, and ATF6 rebalance proteostasis, whereas chronic ER stress leads to terminal UPR [57] that ultimately converges pro-apoptotic signals into apoptotic execution, triggers mitochondrial disruption, and activation of caspase-3 [58]. Reduced cleaved caspase-3 levels in the ONL following VCP inhibition (**Fig. S2**) support this mechanism.

Another factor promoting cell death may involve the high energy cost of ERAD. As an AAA^+^ ATPase, VCP activity consumes energy in the form of ATP equivalents. Thus, reducing VCP activity may alleviate metabolic stress and indirectly improve mitochondrial function [59, 60]. This energy-sparing effect could be particularly beneficial for photoreceptors, given their high metabolic demands.

VCP inhibition has also been demonstrated to increase ER stress, which triggers activation of the UPR. UPR initiates the upregulation of chaperones, including Bip and CHOP, which assist in protein folding [61, 62] and potentially improve folding efficiency. More RHO^WT^ may be released from the ER and trafficked to the plasma membrane following VCP inhibition. Since the antibody used does not discriminate between mutant and WT RHO, this remains speculative and highlights the need for isoform-specific tools.

A recent study has shown that VCP inhibition stabilized misfolded α1(A322D) subunits of the GABA_A_ receptor at the ER [63]. VCP modulation has been shown to ameliorate pathologies of other protein misfolding disorders, including ΔF508-CFTR (which is also a substrate of ERAD) in cystic fibrosis [64, 65] and ninaE*^D^*^1^ in *Drosophila* [17], suggesting broader relevance for neurodegenerative diseases driven by ERAD-sensitive aggregates. These findings highlight the therapeutic potential of VCP inhibition for a range of protein misfolding disorders.

Both VCP inhibitors used here have limited water solubility. Moreover, pharmacokinetic modeling (data not shown) predicts a short intraocular half-life of only a few hours. To overcome these limitations, we encapsulated ML240 in a mPEG-cholane-based nanocarrier [66]. Amphiphilic-based nanocarriers are a new type of nanotechnology-based ODDS that offers numerous advantages over traditional drugs, including enhanced solubility, sustained drug release and facilitated retinal penetration [67, 68]. mPEG-ML240 achieved comparable or superior protection at 2.5 µM compared to 20 µM of free ML240, with only a single intravitreal administration in retinal explants. Compared to free drug and VCP siRNA, mPEG-ML240 more effectively reduced cell death, preserved the ONL, and restored RHO trafficking, likely due to a more sustained delivery at the target site and favorable diffusion characteristics [69, 70]. This supports the feasibility of using nanocarrier-based systems for clinical translation.

Previous studies have shown that small interfering RNA (siRNA)-induced gene silencing is a promising approach for treating chronic eye diseases [32, 33, 35]. Efficient delivery of siRNA across all retinal layers is a huge challenge. Adeno-associated virus (AAV) is considered a suitable vector to introduce exogenous genes into photoreceptors [71–76]; however, immunotoxicity and comparably high costs limit its application [77]. In this study, we used Reverse Magnetofection, a novel, safe, and efficient delivery method, to deliver VCP siRNA across the retina. This non-viral technique achieved gene silencing in the ONL with no detectable toxicity. Similar outcomes were observed using this new ODDS approach in another adRP rat model, *Rho*^P23H^, where VCP expression was inhibited in all retinal layers [41]. Compared to classical magnetofection, the RPE-to-GCL delivery direction in reverse magnetofection provides superior transfection efficiency, consistent with reports from retinal electroporation studies [78]. VCP silencing by Reverse Magnetofection significantly reduced photoreceptor cell death and improved RHO distribution to the OS. Whether this method is suitable for repeated or long-term treatments remains to be determined.

Comparative analysis of the three different VCP inhibitory strategies revealed that mPEG-ML240 nanoparticles exhibited superior efficacy over free ML240 and VCP siRNA in explanted *Rho*^ΔI255/+^ retinae. This enhanced effect could be attributed to the short half-life of both free ML240 and VCP siRNA. Increasing the siRNA dose to 100 nM did not improve the rate of protection and instead reduced efficacy (data not shown). Results from a 16-day intravitreal treatment of *Rho*^ΔI255/+^ mice further confirmed the superior performance of mPEG-ML240. No signs of increased inflammation were observed in retinal explants with any treatment, as indicated by the absence of significant changes in GFAP or Iba1 expression (**Fig. S3**).

*In vivo* OCT imaging showed no significant differences in retinal thickness between eyes treated with mPEG-vehicle and mPEG-ML240 (**Fig. S4**). This is likely due to the fact that OCT measurements are less precise in detecting photoreceptor preservation than histologic analyses. Nevertheless, the preservation of retinal integrity provides important evidence supporting the retinal safety of the treatment over a 16-day period. However, extended safety and efficacy evaluations beyond this time point will be essential to assess the translational potential of each delivery strategy.

In summary, two key findings should be stressed: (1) VCP inhibition confers neuroprotection in a clinically relevant *RHO*^ΔI255/+^ model of adRP, and (2) nanocarrier-based delivery of the hydrophobic VCP inhibitor ML240 (as mPEG-ML240) provides sustained neuroprotection after intravitreal delivery without measurable toxicity. Future studies should explore optimized dosing, long-term efficacy, and potential combination therapies to enhance treatment outcomes further.

## Materials and methods

### Animals

The homozygous *Rho*^ΔI255/ΔI255^ knock-in mouse model was generated by GenOway (Lyon, France), as previously reported [45]. Heterozygous animals (*Rho*^ΔI255/+^) were obtained by crossing homozygous animals with C57BL/6J wild-type (WT) mice (JAX stock #000664). Animals, regardless of gender, were housed in the animal facility of the Tübingen Institute for Ophthalmic Research under standard white cyclic lighting, with *ad libitum* access to food and water. All procedures were conducted in accordance with the Association for Research in Vision and Ophthalmology (ARVO) Statement for the use of animals in ophthalmic and vision research and the German Federal Government (Tierschutzgesetz). Procedures for organ collection used in explant cultures were conducted under internal notifications (Mitteilungen) AK02/20M, AK01/22M, AK08/22M, and AK03/24M. *In vivo* injections were approved under animal experiment license number AK02/23G by the institutional animal welfare office of the University of Tübingen. All efforts were made to minimize the number of animals used and to minimize their suffering.

### Retinal explant cultures of Rho^ΔI255/+^ mice

Retinae were isolated together with the attached RPE as described previously (**Fig. 8A-C**) [44, 79]. Briefly, *Rho*^ΔI255/+^ mice at PN14 and PN17 were sacrificed; the eyes were enucleated under aseptic conditions and pretreated with 12 % proteinase K (MP Biomedicals, 0219350490) for 15 minutes at 37 °C in R16 serum-free culture medium (Invitrogen Life Technologies, 07490743A). The enzymatic digestion was stopped by the addition of 20 % fetal bovine serum (FBS; Sigma-Aldrich, F7524). Retina with attached RPE were dissected, and four radial cuts were made to flatten the tissue. The Tissues were transferred onto a 0.4 μm polycarbonate membrane (Corning Life Sciences, CLS3412), with the RPE side facing the membrane. The inserts were placed into six-well culture plates and incubated in serum-free complete culture medium consisting of Neurobasal^TM^-A (Thermo Fischer Scientific, 10888022), 2% B-27 supplement (Thermo Fischer Scientific, 17504044), 1% N2 supplement (Thermo Fischer Scientific, 17502048), 1% penicillin-streptomycin solution (Thermo Fischer Scientific 15140-122), and 0.4% GlutaMax^TM^ (Thermo Fischer Scientific, 35050061) at 37 °C in a humidified incubator with 5% CO^2^.

Retinae were then randomly assigned to untreated, vehicle, or VCP inhibition groups. The VCP inhibitors, including ML240 (20 μM, TOCRIS, Bio-Techne GmbH, 5153) and NMS-873 (0.5 μM and 1 μM, Xcess Biosciences, M60165-b), dissolved in DMSO (Sigma-Aldrich, D2650), were applied to the culture medium at PN14 and again with a change of medium every two days until PN20, the peak of degeneration in age-matched *in vivo* mutants. ML240 and NMS-873 were dissolved in DMSO to obtain a stock concentration of 1 mM each. The 1 mM stock was used to prepare the final therapeutic dose in Neurobasal^TM^-A medium. DMSO was added where necessary to always achieve a final DMSO concentration of 0.5%. Drugs were sonicated (Sonorex Super RK 103H, Bandelin) for 1 min, 3 times at room temperature to obtain a homogeneous formulation. DMSO (0.5%) alone, diluted in the culture medium, was used to generate the corresponding vehicle control. mPEG-cholane was synthesized according to previously published protocols (**Fig. 8D**) [27], and ML240-loaded nanoparticles were generated based on the ‘film hydration’ method as previously described [66]. The mPEG-cholane encapsulated ML240 (2.5 μM) was administered only once on top of the GCL side of the retina at PN14 (**Fig. 8F-b**), followed by medium replacement until PN20. Drug-free mPEG-cholane at equivalent concentrations was used as the vehicle control. In both drug delivery systems, untreated retinae with Neurobasal^TM^-A medium were used as controls.

### Reverse Magnetofection of VCP siRNA in Rho^ΔI255/+^ retinal explants

MNPs (XPMag) were purchased from OZ Bioscience (Marseille, France). All siRNAs were non-modified 21-mer with double single-stranded RNA bases in 3’. VCP siRNA (mouse; Thermo Fischer Scientific, Ambion, 152438) and scrambled siRNA (Thermo Fischer Scientific, Ambion, 4390843; negative control) were used in this study. This experiment (**Fig. 8E**) was carried out as described before [41]. Briefly, XPMag/ siRNA complexes were prepared by mixing 2.5 µL of XPMag with 50 nM VCP siRNA or scrambled siRNA, followed by incubation for 30 min at room temperature in 100 µL of serum- and complement-free Neurobasal^TM^-A medium. Afterward, magnetic complexes were added to 900 µL of complete Neurobasal^TM^-A medium preplaced in six-well culture plates. Reverse Magnetofection was applied by placing the super-magnetic plate above the lid of the culture plate. After 30 min, the magnetic field was removed, and retinal explants were incubated under standard conditions (**Fig. 8F-c**). Twenty-four hours later, the culture medium was replaced with fresh medium for another 48 h before *VCP* silencing evaluation was performed. Untreated retinae, retinae treated with scrambled siRNA, and retinae treated with XPMag alone (no siRNA) were used as controls. *Rho*^ΔI255/+^ retinae were treated at PN17 and collected for retinal cryosection on PN20.

### Fixation and preparation of retinal cryosections

Retinal explants from *Rho*^ΔI255/+^ mice were fixed in 4% paraformaldehyde (PFA, Polysciences, Inc.) in 0.1 M phosphate buffer (PB; pH 7.4) for 45 min at 4 °C, followed by cryoprotection in graded sucrose solutions (10% for 10 min, 20% for 20 min, 30% for 1h) and embedded in Tissue-Tek^®^ OCT Compound (Sakura^®^ Finetek, VWR, 4583). Vertical sections (14 µm thick) were collected, air-dried, and stored at -20 °C until use.

### TUNEL assay

TUNEL (terminal deoxynucleotidyl transferase dUTP nick end labeling) assay was performed using an *in situ* cell death detection kit conjugated with fluorescein isothiocyanate (Roche, 11684795910). DAPI (Sigma, D9542) was used as a nuclear counterstain.

### Immunohistochemistry

Immunofluorescence staining was performed on cryosections from *Rho*^ΔI255/+^ retinal explants. Retinal Sections were incubated overnight at 4 °C with the primary antibodies (RHO: Sigma-Aldrich, MAB5316, 1:350; VCP: Invitrogen, MA3-004, 1:1000; GFAP: Merck Millipore, G3893, 1:500; Iba1: Fujifilm Wako Chemicals, 019-19741, 1:400; caspase-3: Cell signaling, 9664, 1:300). Fluorescence immunocytochemistry was performed using Alexa Fluor® 568 (goat anti-mouse IgG: Thermo Fischer Scientific, A-11031, 1:500; goat anti-rabbit IgG: Thermo Fischer Scientific, A-11036, 1:500) or Alexa Fluor® 488 (goat anti-mouse IgG: Thermo Fischer Scientific, A-11029, 1:500) conjugated secondary antibody. Negative controls were included by omitting the primary antibody.

### Scanning-laser ophthalmoscopy (SLO) and optical coherence tomography (OCT)

*In vivo* imaging was performed as previously described [80, 81]. Animals were examined immediately after anesthesia at PN30. A 100 diopter (dpt) contact lens was placed on the cornea after applying hydroxypropyl methylcellulose to avoid dehydration. SLO imaging was performed in conjunction with OCT on the Spectralis™ HRA+OCT device (Heidelberg Engineering, Heidelberg, Germany) equipped with a superluminescent diode at 870 nm as low the coherence light source and operated via the proprietary software Eye Explorer version 5.3.3.0. Each two-dimensional B-scan, recorded with a 30° field of view, contained up to 1536 A-scans and was acquired at a rate of 40,000 scans per second. Data were exported as 8-bit greyscale images. For retinal thickness measurements, reflectivity profiles were extracted between the ganglion cell layer and the retinal pigmented epithelium at three distinct points within the OCT scan, as previously described [82].

### Microscopy and image analysis

All immunostaining samples were analyzed using a Zeiss Axio Imager Z1 ApoTome microscope, AxioCam MRm camera, and Zeiss Zen 2.3 software in Z-stack at 20x magnification. TUNEL-positive cells in the ONL of at least three sections per group were manually counted for quantitative analysis. The percentage of TUNEL-positive cells and activated caspase-3 cells was calculated by dividing the number of positive cells by the total number of ONL cells. Photoreceptor cell rows were assessed by counting the individual nuclei rows in one ONL and averaging the counts.

The intensity mean value of RHO immunofluorescence was measured by contouring the IS together with the ONL and the OS areas. To quantify changes in RHO distribution, the ratio of RHO mean fluorescence intensity (MFI) was calculated using the following formula:

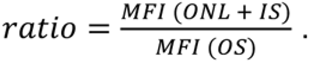

For VCP, mean immunofluorescence intensity was measured by outlining the ONL region using Zeiss Zen software..OS thickness was measured via the distance tool in Zeiss Zen software.

### Intravitreal injection of Rho^ΔI255/+^ mice

*Rho*^ΔI255/+^ mice of at least 5 g body weight were treated at PN 14. Animals were anesthetized with a subcutaneous injection of ketamine (66.7 mg/kg) mixed with xylazine (11.7 mg/kg) and diluted in 0.9% NaCl saline. Tropicamide drops (Pharma Stulln, Stulln, Germany) were applied to each eye for pupil dilation. A Hamilton syringe (Hamilton Robotics, Bonaduz, Switzerland) with a 33-gauge (G) needle was used to intravitreally inject a single 2 μl volume of a sterile preparation of mPEG-ML240 into the left or right eye or an equivalent volume of mPEG-vehicle (empty) into the other eye. Tobramycin (Tobrex) was applied to the eye after injection to protect the cornea and prevent infection. Animals were then euthanized at PN30 and eyeballs were collected for histological staining. To distinguish between the superior and inferior hemispheres of the retina, red dye was nasally marked onto the eyeball before fixation with 4% PFA. Subsequently, fixed eyeballs were dehydrated, embedded, and stained with TUNEL assay and RHO antibody, as described above.

### Statistics

Statistical analysis was performed using GraphPad Prism 10.2.1 software. A one-way ANOVA was used to evaluate differences in the percentage of TUNEL-positive cells in the ONL, number of ONL rows, RHO mean FL intensity, OS length, and the comparison of VCP inhibition neuroprotective effects across three different inhibitory formulations in retinal explants. When a 0.05 level of significance was found, Tukey post-hoc tests were performed to compare different groups. For intravitreal injection experiments, retinae treated with mPEG-vehicle and mPEG-ML240 were compared using a paired two-tailed *t*-test. All the above-mentioned data were presented as box-and-whisker plots (median, quartiles, min-max). An unpaired two-tailed t-test was applied to analyze the relative percentage of caspase-3-positive cells compared with the untreated control group in three different VCP inhibitory delivery systems. Quantitative data are shown as the mean ± SD; *p* < 0.05 was considered statistically significant.

## Authorship contribution statement

**Bowen Cao**: Conceptualization, Methodology, Validation, Formal analysis, Investigation, Data curation, Writing-original draft, Writing-review and editing, Visualization. **Regine Mühlfriedel**: Methodology, Investigation. **Merve Sen:** Methodology, Investigation. **Ana-Cristina Almansa-**

**Garcia:** Methodology, Investigation. **Mathias W. Seeliger**: Methodology, Investigation, Data curation, Writing – review. **Anne-Sophie Petremann-Dumé**: Methodology, Validation, Investigation. **Ellen Kilger**: Conceptualization, Methodology, Writing-review and editing, Supervision, Project administration. **Anneli Vollert**: Methodology, Investigation. **Sylvia Bolz**: Methodology, Investigation. **Christine Henes:** Methodology, Investigation. **Paolo Caliceti**: Methodology, Writing - review. **Stefano Salmaso**: Methodology, Writing - review. **Marius Ueffing**: Conceptualization, Validation, Formal analysis, Investigation, Resources, Writing-review and editing, Supervision, Project administration, Funding acquisition. **Blanca Arango-Gonzalez**: Conceptualization, Validation, Formal analysis, Investigation, Data curation, Writing-review and editing, Supervision, Project administration, Funding acquisition.

## Acknowledgments

This research was supported by the Tistou and Charlotte Kerstan Foundation (www.kerstanstiftung.org; RHO-Cure program); Fighting Blindness Canada (FBC, www.fightingblindness.ca; Transformative Research Award: - TargetVCP); ProRetina foundation (www.pro-retina-stiftung.de); Zinke Heritage foundation (awarded to MUe), and the China Scholarship Council (CSC). The animal husbandry personnel at the Universitätsklinikums Tübingen and Norman Rieger are acknowledged for their animal care.

