## Supplementary Material for "Ocular delivery of different VCP inhibitory formulations prevents retinal degeneration in rhodopsin 255 isoleucine deletion mice"

* *Equal last authorship*

### Supplementary Materials

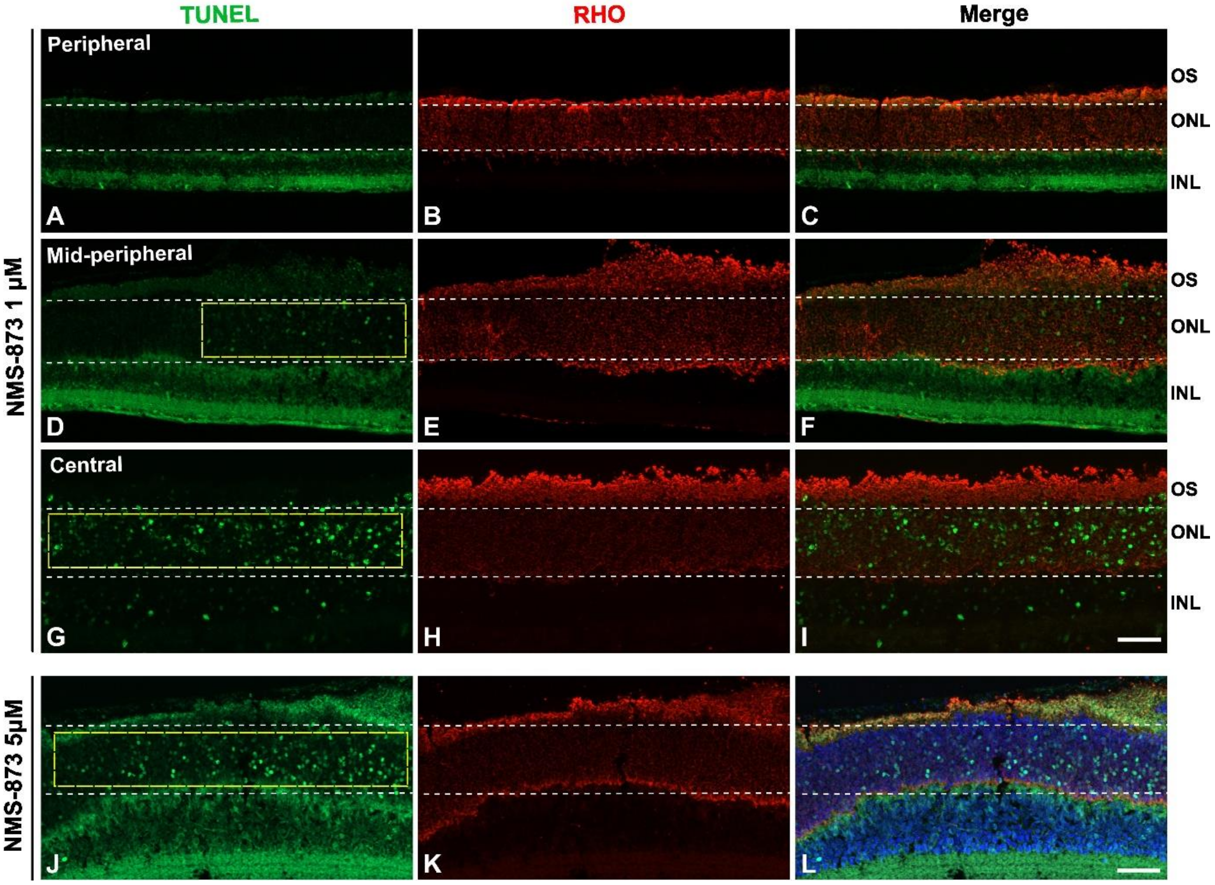


**Figure S1. Toxicity of different doses of NMS-873 in different regions of *Rho*$^{\Delta I255/+}$ retinal explants.** **R**etinae treated with 1 µM NMS-873 showed a reduction in TUNEL-positive cells in peripheral retinae (**A**) and predominant localization of RHO to the OS (**B**). However, an abnormal increase in TUNEL-positive cells throughout the ONL was observed in the mid-peripheral (**D**, yellow dashed box) and central retinae (**G**, yellow dashed box) was observed. In addition, after treatment with a higher concentration of 5µM NMS-873, widespread cell death was detected, with TUNEL-positive cells filling the entire ONL (**J**, yellow dashed box). Interestingly, these two different doses of NMS-873 improved the trafficking of RHO to the OS (**B**, **E**, **H**, **K**). Merged images of TUNEL and RHO staining are shown in **C**, **F**, **I**, and **L**. Scale bar: 50 µm. *Rho:* rhodopsin gene, RHO: rhodopsin protein.

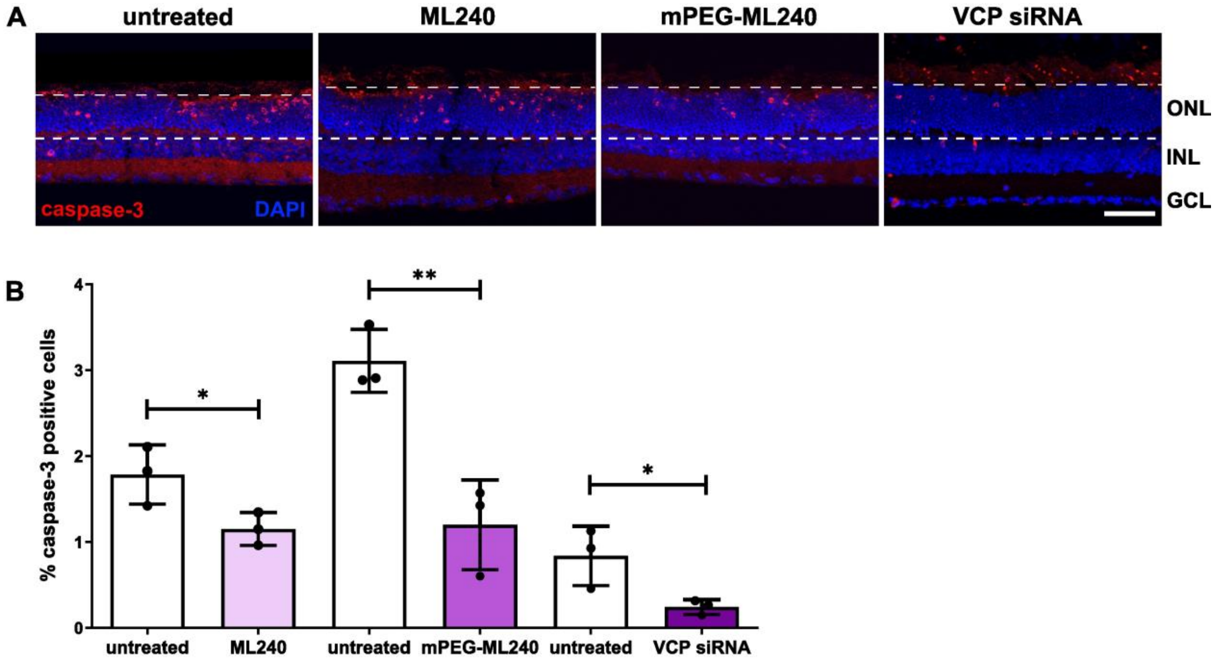


**Figure S2. Different VCP inhibitory formulations reduce activated caspase-3 levels in *Rho*$^{\Delta I255/+}$ retinae. (A)** Caspase-3-positive cells in the retinal ONL were stained with an antibody against cleaved caspase-3 (red). Nuclei were counterstained with DAPI (blue). Scale bar: 50 µm. **(B)** Quantification of the percentage of caspase-3-positive cells relative to the total number of ONL cells. VCP inhibition led to a significant reduction in caspase-3-positive cells compared to untreated retinae. Values were quantified from 3 retinae (black dots). Data presented as mean ± SD; Statistical analysis: Unpaired t-test; *$p < 0.05$, **$p < 0.01$.

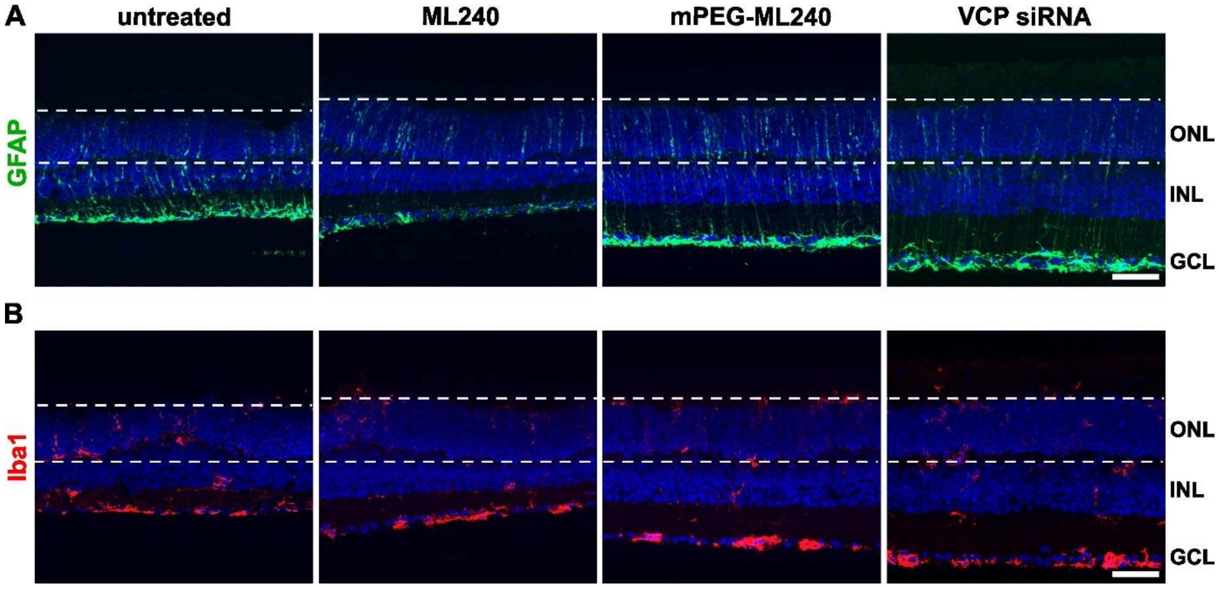


**Figure S3. Different VCP inhibitory formulations and drug delivery systems do not aggravate associated retinal gliosis or microglial migration in *Rho*$^{\Delta I255/+}$ mice**. Immunofluorescence labeling of retinal cryosections from untreated and VCP inhibitor-treated retinal explants showed the location of activated Müller cells, indicated by GFAP immunoreactivity (**A**, green) and microglial cells, indicated by Iba1 staining (**B**, red). Nuclei were counterstained with DAPI (blue). The application of different forms of VCP inhibition neither aggravated nor alleviated inflammatory responses compared to the untreated group. Scale bar: 50 µm. GFAP: glial fibrillary acidic protein, Iba1: ionized calcium-binding adapter molecule 1.

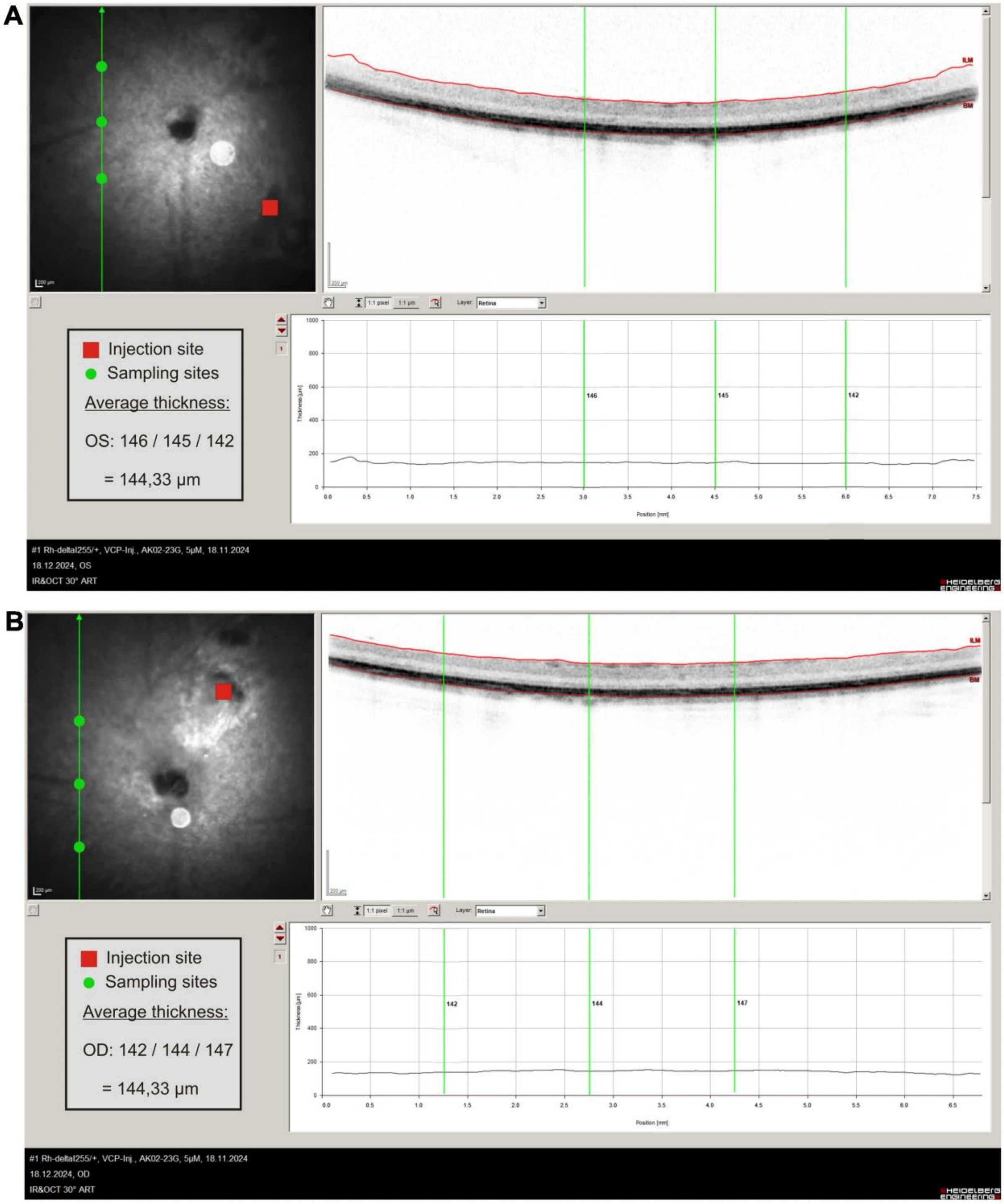


**Figure S4. *In vivo* fundus image and retinal morphology of *Rho*$^{\Delta I255/+}$ mice after long-term treatment**. *Rho*$^{\Delta I255/+}$ mice treated with 5 µM mPEG-vehicle (**A,** left eye, OS) and mPEG-encapsulated ML240 (**B,** right eye, OD) for 16 days showed no differences in fundus images determined by scanning laser ophthalmoscopy (SLO), nor in retinal thickness measured by OCT, suggesting that prolonged VCP inhibition does not induce toxicity in the retinae.

.
